# Acute lysosome repositioning reveals functional and proteomic adaptations

**DOI:** 10.64898/2026.05.17.724935

**Authors:** V. Dostál, A. R. Pollio, S. Kofler, C. Krebiehl, L. Kremser, B. Sarg, T. Stasyk, L. A. Huber

## Abstract

Lysosomes exhibit spatial heterogeneity, but establishing causal relationships remains challenging with existing relocalization strategies. We present a modular toolkit that rapidly repositions lysosomes on demand by recruiting inducible motors. This decouples the location of organelles from systemic stress. We demonstrate that peripheral lysosomes adapt quickly by exhibiting luminal alkalinization and reduced proteolytic capacity. Furthermore, peripheral sequestration impairs autophagic flux by creating a spatial “trafficking bottleneck”. We use this system to provide an unbiased proteomic characterization of spatially distinct lysosomal populations using mass spectrometry. Our findings reveal a distinct set of proteins and complexes that are spatially partitioned between perinuclear and peripheral lysosomes. Perinuclear lysosomes are configured for metabolic recycling. They have a high density of V_1_ V-ATPase subunits and contain the nucleoside transporter SLC29A3. Conversely, peripheral lysosomes serve as secretory outposts that are enriched in cathepsin Z and the SPG11/SPG15/AP5 complex. These validated cell lines and extensive datasets provide a flexible framework for investigating the functional specialization of different lysosome populations.

## Introduction

Lysosomes (or, more broadly, endolysosomes) are ubiquitous cellular organelles which are increasingly recognized for their functional versatility. Besides their well-known role in the degradation of cellular waste and nutrient recovery, endolysosomes also participate in cell signaling and homeostatic control (Sancak et al., 2010; Settembre et al., 2012), lipid and calcium exchange (Infante et al., 2008; Shen et al., 2012), secretion into the cellular exterior, plasma membrane repair (Reddy et al., 2001; Jaiswal et al., 2002) and immunity (Gui et al., 2019). These functions are enabled by more than 300 endolysosomal proteins and at least the same number of additional proteins which interact with lysosomes in some way (Mosen et al., 2021); more lysosomal proteins are discovered every year.

Rather than existing in a uniform state, lysosomes exhibit significant heterogeneity, pointing toward a model of spatial and functional specialization (Chavan and Bhattacharjee, 2025). Lysosomes widely differ in their size, motility, ion content and their position within the cytoplasm (Cabukusta and Neefjes, 2018; de Araujo et al., 2020; Halcrow et al., 2022). Individual endolysosomes display distinct marker behavior, with the presence or absence of well-known markers TMEM192 or NPC1 suggesting diversification (Bond et al., 2025). However, identifying lysosomal subpopulations with unique proteomic signatures remains challenging due to a lack of high-resolution tools. While methods like HA-tag-assisted native immunoprecipitation (LysoIP) or loading endolysosomes with paramagnetic iron (FeDex) effectively isolate lysosomes in bulk (Singh et al., 2020), they lack the spatial precision required to resolve lysosomal heterogeneity.

Endolysosomes move between the perinuclear region and the cell periphery in response to various stimuli, such as starvation (Korolchuk et al., 2011) or stress (Sasazawa et al., 2022). Anterograde (kinesin-directed) movement of endolysosomes is mediated by Arl8 and PLEKHM2 in conjunction with the BORC complex (Rosa-Ferreira and Munro, 2011; Pu et al., 2015) or via an alternative pathway instrumented by FYCO1 and protrudin (Hong et al., 2017). Conversely, dyneins are recruited for retrograde trafficking primarily by RILP or via alternative pathways involving TMEM55B, JIP4 and other proteins (Jordens et al., 2001; Li et al., 2016; Willett et al., 2017). The diversity of regulatory pathways controlling lysosomal trafficking highlights the fundamental importance of organelle positioning in maintaining cellular homeostasis.

The spatial positioning of lysosomes dictates their structural and functional identity. Compared to their perinuclear counterparts, peripheral lysosomes exhibit reduced acidity and enzymatic activity (Johnson et al., 2016). This is likely due to a twofold reduction in V-ATPase complex density per vesicle (Maxson et al., 2022). Their proximity to the plasma membrane facilitates frequent exocytosis (Jaiswal et al., 2002) ‒ a process that can degrade the extracellular matrix and increase melanoma cell invasiveness (Jerabkova-Roda et al., 2025). Additionally, these populations have different “GTPase blueprints”: perinuclear lysosomes are enriched in Rab7, while peripheral lysosomes preferentially recruit Arl8b (Johnson et al., 2016; Schleinitz et al., 2023).

These spatial differences define the identity of lysosomal pools. However, determining their global physiological impact remains a significant challenge. A prime example is the regulation of mTORC1 signaling. Some studies suggest that peripheral repositioning enhances mTORC1 activity (Korolchuk et al., 2011), while others have observed the opposite effect (Clippinger and Alwine, 2012; Walton et al., 2019). These discrepancies likely stem from the various and sometimes invasive techniques employed to manipulate lysosome position, such as dynein inhibition or medium acidification, which can introduce secondary effects that complicate interpretation of the results (Ballabio and Bonifacino, 2020).

Lysosomal position also likely regulates the frequency of encounters between autophagosomes and lysosomes, thus playing a role in autolysosome formation. The specific outcome of repositioning is again not completely clear, depending on the preferred cellular site of autophagosome-lysosome fusion: there are reports that this mainly happens in the periphery (Jia et al., 2017) or that autophagosomes are rather delivered for clearance to the perinuclear region (Cardoso et al., 2009; Korolchuk et al., 2011). Clearly, responses to lysosomal repositioning are strongly context- and cell type-dependent. However, our understanding of this diversity has been hindered by a lack of tools that can isolate positioning from off-target effects. Here, we use an inducible FKBP/FRB* recruitment system to bypass traditional techniques and rapidly relocalize lysosomes via motor-driven processes.

Our results conclusively demonstrate that endolysosomes are not static, but rather acutely adapt their functional identity to their intracellular niche. Additionally, we use this system to create the first comprehensive proteomic map of perinuclear versus peripheral lysosomes. This reveals the molecular signatures that drive organelle specialization.

## Results

### Development and validation of inducible cellular models for acute lysosomal repositioning

We developed FKBP/FRB* stable cell lines that allow rapid repositioning of (endo)lysosomes to the perinuclear region or to the periphery in response to A/C (AP21967) treatment (Bayle et al., 2006). To ensure the broad applicability of our findings, we used a panel of three complementary parental cell lines: U2OS, a human osteosarcoma line that is a well-established model for lysosomal dynamics; hTERT RPE-1, an immortalized human retinal pigment epithelial line that is favored for its diploid genome and organized architecture, which makes it ideal for high-resolution microscopy and quantitative proteomics; and mouse embryonic fibroblasts (MEFs), which provide a primary-like physiological context for validating our observations across species and lineages.

The repositioning system consists of two distinct modules (**Figure 1 a**). First, cells were transduced with a lysosomal-targeting “anchor” comprising the 39 N-terminal amino acids of LAMTOR1 (LT1 tag) fused to mCherry and an FRB* (T2098L) domain (Nada et al., 2009; Amick et al., 2019). We utilized the LT1 tag for anchoring FRB* to ensure a more physiological targeting mechanism, as recent studies have indicated that other anchors including LAMP1 overexpression can perturb endogenous lysosomal function (Shah et al., 2023; Zhang et al., 2023; Cheetham-Wilkinson et al., 2025). In a second step, these cells were transduced with FKBP fused to either BicD2^2–594^ or to KIF5C^1–559^, which mediate (-)- or (+)-end directed microtubule transport, respectively (Bentley et al., 2015). These constructs additionally contained 3×HA tags at their N-termini to facilitate biochemical studies.

**Figure 1:**
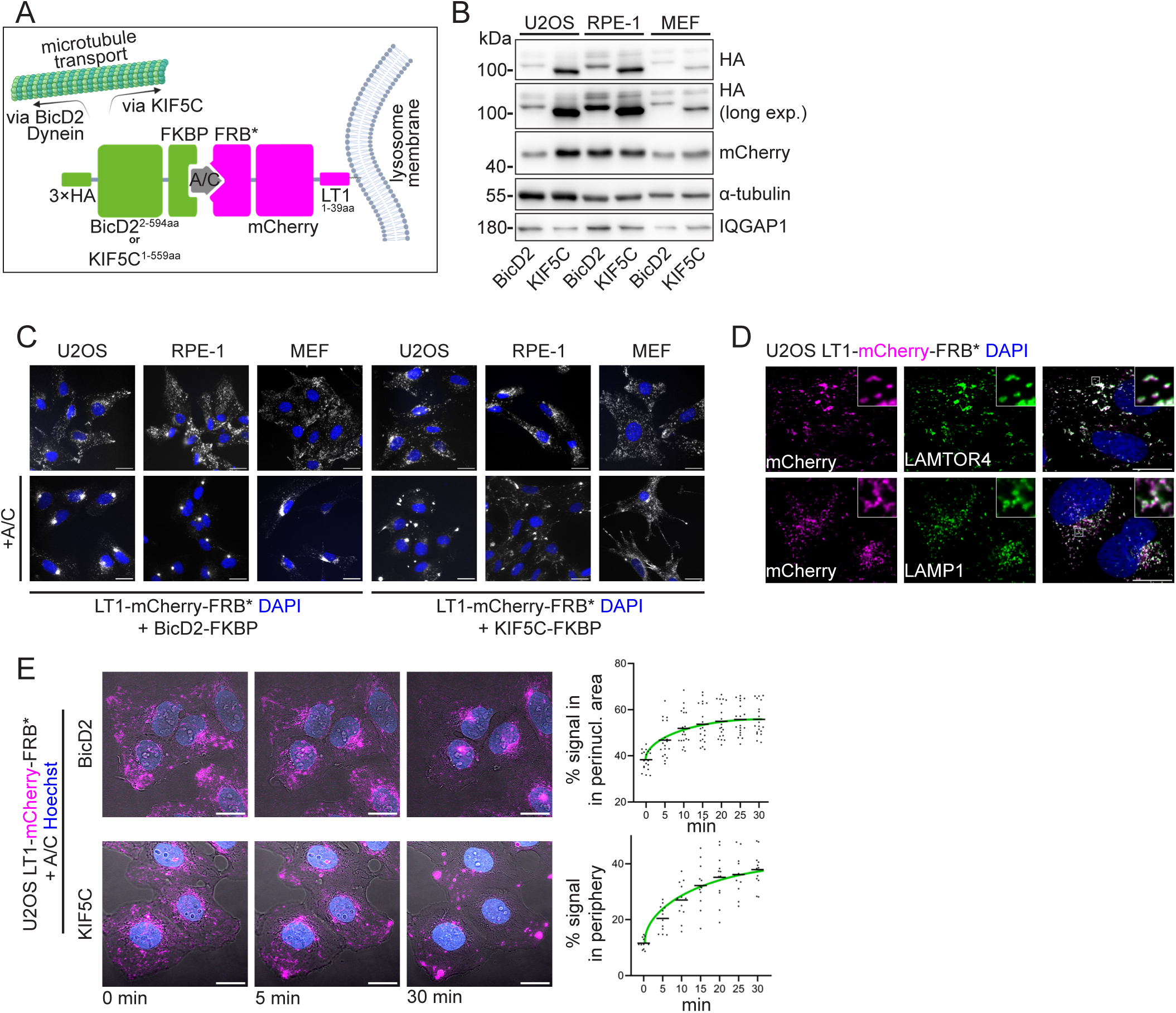
Development and validation of the lysosome repositioning cell lines. (A) Schematic diagram depicting the design of the repositioning cell lines. (B) Western blot showing expression of lysosomally targeted mCherry-FRB* protein and a 3×HA tagged FKBP fused to BicD2 or KIF5C, each in three different parental cell lines. (C) Microscopical confirmation of the repositioning of mCherry-tagged lysosomes (grey) in each of the cell lines. Cells were treated with A/C or control, fixed and mCherry localization was recorded by epifluorescence. Nuclei in blue (DAPI). Scale bar: 25 μm. (D) Airy-scan confocal microscopy of U2OS cell lines expressing the LT1-mCherry-FRB* protein (magenta), stained for LAMTOR4 or LAMP1 (green) and nuclei (blue). Scale bar: 15 μm. (E) Left: Live imaging of U2OS repositioning cell lines showing a brightfield view overlaid with mCherry (magenta) and a nuclear Hoechst stain (blue). Scale bar: 20 μm. See also **Video 1**. Right: Quantification of % mCherry signal in the 20% most perinuclear or in the 20% most peripheral area of cells. Logistic curves (green) were fitted to the data points. Each data point = 1 cell.

Stable clones for all three parental lines (U2OS, hTERT RPE-1, and MEF) successfully expressed both the lysosomal anchor and the motor fusion proteins, although we noted that BicD2 expression was consistently lower than that of KIF5C (**Figure 1 b**). While non-induced cell lines exhibited minor baseline differences in distribution ‒ with non-induced BicD2 lines appearing more peripheral than non-induced KIF5C lines ‒ the addition of the A/C heterodimerizer effectively mobilized lysosomes across the entire population toward their respective target region. The repositioning response was most robust in U2OS and hTERT RPE-1 cells, whereas MEFs exhibited a more modest redistribution (**Figure 1 c**). Consequently, we focused our functional and proteomic analyses on the human U2OS and hTERT RPE-1 (henceforth “RPE-1”) cell lines.

To verify the precision with which the repositioning modules target, we characterized the localization of the LT1-mCherry-FRB* anchor relative to endogenous lysosomal markers. Using super-resolution (Airyscan) microscopy on U2OS cells, we observed nearly perfect colocalization between the LT1-driven construct and LAMTOR4. This result confirms that the LT1 tag accurately directs the system to the Ragulator-associated lysosomal microdomain (**Figure 1 d**). Furthermore, although the LT1 construct showed significant overlap with LAMP1, super-resolution imaging revealed that these markers frequently occupy separate but adjacent microdomains on the same endolysosomal vesicle. This high-resolution spatial segregation aligns with the lysosomal membrane’s known modularity. Together, these data demonstrate that our system specifically targets a well-defined population of LAMP1-positive endolysosomes by utilizing their endogenous LAMTOR-associated scaffolding machinery.

We then determined the optimal concentration of A/C to be used for repositioning of lysosomes in our cell lines, and the overall dynamics of this process. First, the U2OS repositioning cell lines were induced with a range of A/C concentrations, fixed, and stained with an anti-HA antibody. We observed relocalization of mCherry signal to either perinuclear or peripheral patches and concurrent mobilization of a fraction of HA signal to these spots, reflecting FRB*-FKBP binding. We determined that A/C concentrations equal or greater than 5 nM reliably induce repositioning throughout the whole cell population (**Figure S1 a**), which is in agreement with the reported EC_50_<10 nM (Liberles et al., 1997).

Because the A/C heterodimerizer is a rapamycin analog, we first determined an appropriate working concentration below the level of mTORC1 inhibition to avoid confounding metabolic effects. Western blot analysis confirmed that 5nM A/C ‒ the dose used for all subsequent experiments ‒ does not alter the phosphorylation levels of p70 S6 kinase (S6K), a canonical downstream target of mTORC1. However, we only observed significant mTORC1 inhibition at concentrations ≥50nM (**Figure S1 b-c**). The dynamic response to 5nM A/C treatment is remarkably rapid, with lysosomal mobilization detectable within 5 minutes, and near-complete redistribution achieved by 30 minutes (**Figure 1 e** and **Video 1**). Importantly, this acute repositioning does not compromise cellular homeostasis; cells remain fully viable, and their proliferation rates are indistinguishable from non-induced controls. After division, lysosomes in daughter cells retain their localization. The process is slowly reversible upon drug washout, with approximately 50 % of the cell population returning to a baseline lysosomal distribution after 48 h (**Figure S1 d-f**). These results suggest that inducing lysosomal transport does not disrupt microtubule-based traffic or cellular metabolism globally. This provides an efficient, rapid and safe protocol for lysosomal repositioning.

### Peripheral lysosomes undergo acute luminal alkalinization and show reduced proteolytic activity

To determine if repositioned endolysosomes adapt to their new intracellular niches, we first measured their luminal pH using the ratiometric indicator LysoSensor Yellow/Blue Dextran. After 4 h of A/C treatment, a duration hypothesized to allow metabolic adaptation, we observed that peripheral lysosomes were significantly less acidic than their perinuclear counterparts in both the U2OS and RPE-1 cell lines (**Figure 2 a**). The average pH is within the range reported for these or similar cells (Webb et al., 2021; Rennick et al., 2022; Deng et al., 2025), confirming that endolysosomes acutely tune their luminal environment in response to spatial positioning.

**Figure 2:**
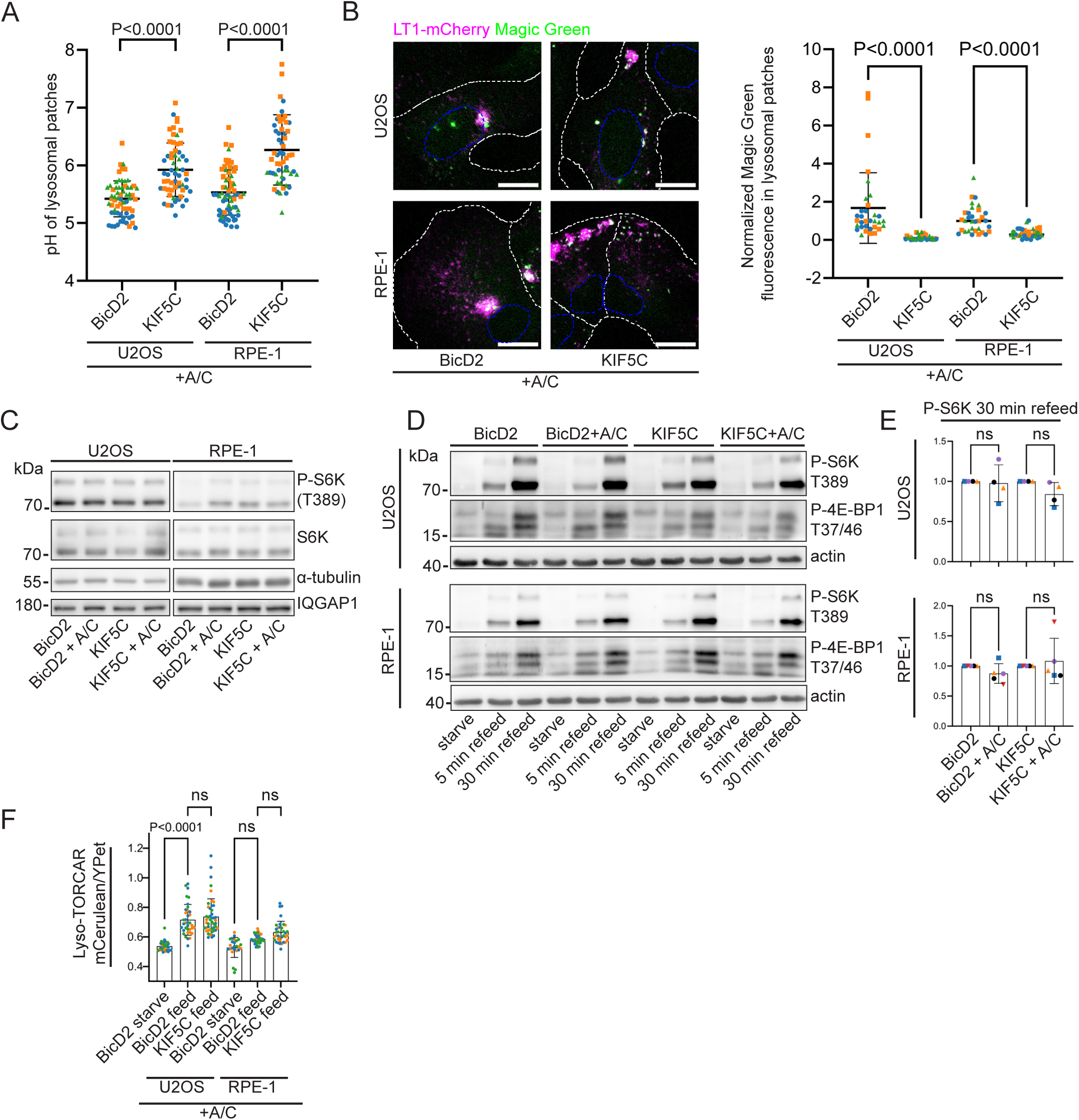
Acidity, hydrolytic activity and mTORC1 signaling competence of repositioned lysosomes. (A) Measurement of lysosomal pH in U2OS and RPE-1 repositioning cell lines fed with LysoSensor Yellow/Blue Dextran and imaged live after A/C induction (4 h). N=3 independent replicates. Each data point = 1 lysosomal patch. Lines: Mean +/- SD. Kruskal-Wallis test for data with non-normal distribution, followed by Dunn’s multiple comparisons test were used. (B) Left: Confocal image of Magic Green fluorescence in U2OS and RPE-1 repositioning cell lines induced with A/C (4 h), showing mCherry (magenta) and Magic Green (green). Outlines of cells and nuclei are shown in white and blue, respectively. Scale bar: 15 μm. Right: Quantification of Magic Green intensity in the lysosomal patches. Values were normalized to Dextran-A647 fluorescence in each lysosomal patch and to median BICD2 sample intensity in each replicate. N=3 independent replicates. Each data point = 1 lysosomal patch. Lines: Mean +/- SD. Kruskal-Wallis test for data with non-normal distribution, followed by Dunn’s multiple comparisons test were used. (C) Western blot showing S6K phosphorylation levels as a measure of steady state mTORC1 activity in the repositioning cell lines with or without A/C (4 h). (D) Western blot showing S6K and 4EBP1 phosphorylation levels in starved cells and upon 5min or 30min restimulation, with or without A/C (4 h). (E) Densitometric quantification of S6K phosphorylation levels from Western blot data (30 min restimulation). N=4 independent replicates, colors of data points indicate the replicates. Ordinary one-way ANOVA was used. Lines: Mean +/- SD. (F) Quantitative FRET assay for lysosomal mTORC1 activity using Lyso-TORCAR. Each data point = 1 lysosomal patch. N=3 independent replicates. Lines: Mean +/- SD. Kruskal-Wallis test for data with non-normal distribution, followed by Dunn’s multiple comparisons test were used.

Next, we reasoned that deacidification should diminish degradative capacity by shifting the luminal pH away from the optimum of acid hydrolases. Although total cellular proteolysis, as measured by DQ Ovalbumin, remained unchanged upon repositioning (**Figure S2 b-c**), high-resolution analysis of individual lysosomal patches revealed a localized effect. By measuring fluorogenic cathepsin substrate (Magic Green) normalized to fluorescent dextran (Johnson et al., 2016), we found that the peripheral endolysosomes are significantly less proteolytically active than the perinuclear lysosomes (**Figure 2 b** and **S2 a**). These results indicate that while global degradation may be compensated, proteolytic efficiency is spatially compartmentalized, with the highest activity concentrated in the perinuclear region.

### Lysosomal positioning does not dictate mTORC1 signaling competence

Conflicting reports have suggested that mTORC1 signaling preferentially occurs at either the perinuclear or peripheral lysosomal pools (Korolchuk et al., 2011; Walton et al., 2019). However, we could not observe any changes in the phosphorylation of canonical targets of mTORC1 in fed conditions or in a starved/refed setup (**Figure 2 c–e**) upon induced relocalization of lysosomes. We then hypothesized that the signaling differences may only be measurable at the level of individual repositioned lysosomal patches. We employed the Lyso-TORCAR system which uses a pair of fluorescent proteins with a linker made of 4E-BP1 to measure FRET change as a result of mTORC1-dependent 4E-BP1 phosphorylation (Zhou et al., 2015). The probe suffers from a lower dynamic range; starvation significantly decreases FRET ratio (mCerulean/YPet) in U2OS but not in RPE-1 cells. Nevertheless, we did not observe any differences in the FRET ratio between perinuclear and peripheral lysosomes in U2OS or RPE-1 cells (**Figure 2 f**). We conclude that the induced location of endolysosomes in U2OS and RPE-1 cells does not affect their ability to serve as signaling platforms for mTORC1.

### Peripheral positioning promotes constitutive lysosomal exocytosis in RPE-1 cells

Lysosomal exocytosis is another key function attributed to the peripheral lysosomes (Jaiswal et al., 2002). We assayed lysosomal exocytosis by measuring beta-hexosaminidase in the medium of repositioned cell lines. After confirming reliability of this assay with ionomycin treatment, which greatly increases the release of beta-hexosaminidase into the medium of U2OS and RPE-1 cells, we similarly measured constitutive exocytosis in the repositioning cell lines over a 24h time window. Although we observed no differences in U2OS cells, which generally show lower levels of lysosomal exocytosis, we confirmed a significant increase in RPE-1 cells with lysosomes repositioned to the periphery (**Figure S2 d-e**). These findings confirm that proximity to the plasma membrane is a key determinant for exocytic competence, particularly in epithelial cell models like RPE-1.

### The spatial segregation of lysosomes impairs the clearance of autophagy

Finally, we examined the influence of lysosomal positioning on the final stages of autophagy, specifically autophagosome-lysosome fusion and subsequent autolysosome clearance. To induce autophagy, we starved U2OS and RPE-1 cells in an amino acid- and serum-free medium for 2 h. Then, we performed immunostaining for the autophagosomal marker LC3.

Starvation led to the formation of LC3-decorated puncta representing autophagosomes and their different stages of maturation. We first measured the overlap of these puncta with mCherry to quantify how autophagosomes merge with lysosomes with or without A/C present in the medium. Manders coefficient for the overlap of LC3 with mCherry was reduced in cells with peripheral lysosomes, indicating problems with autophagosome-lysosome fusion in these cells (**Figure 3 a–b**).

**Figure 3:**
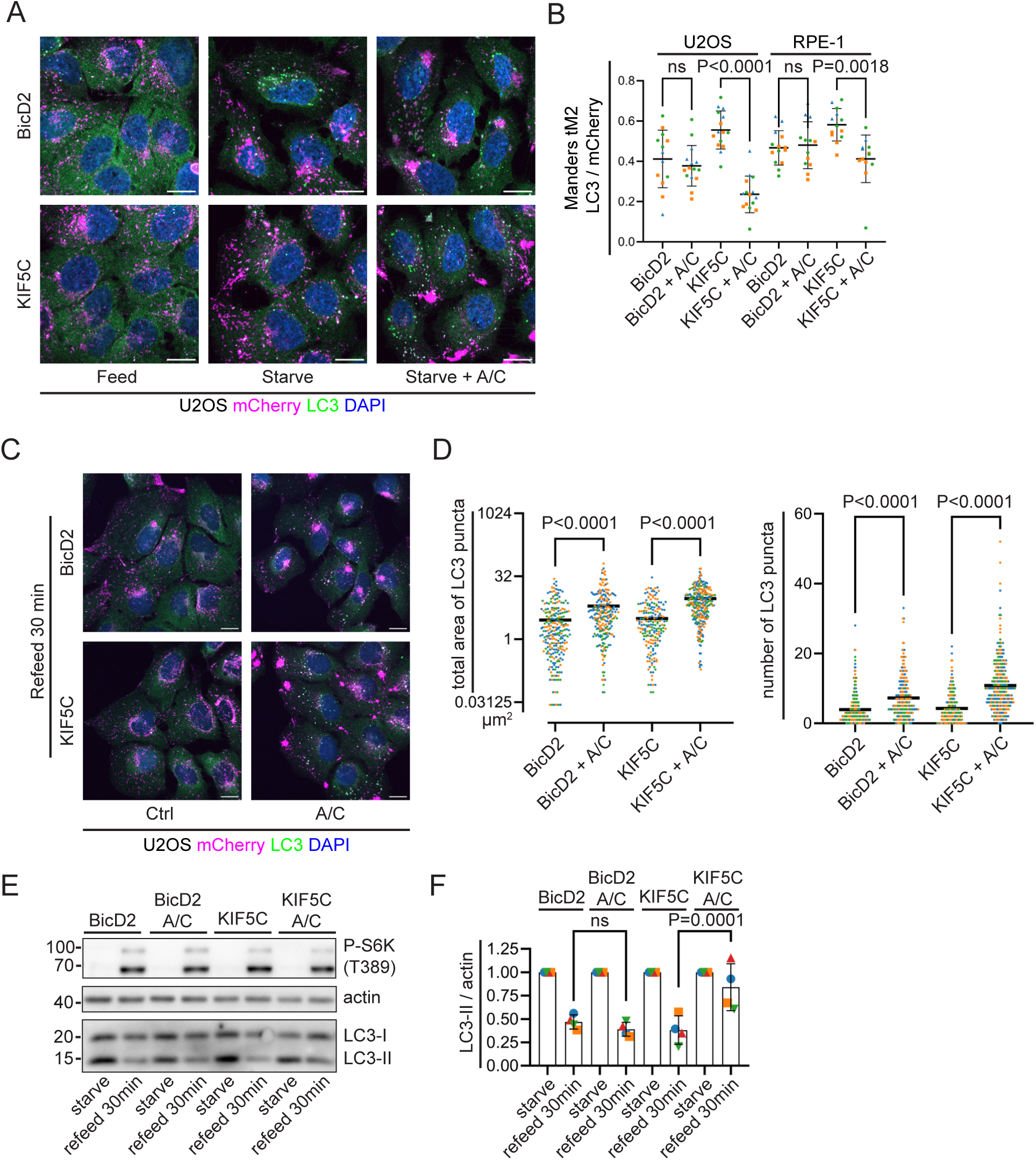
Autophagic flux in the lysosome repositioning cell lines. (A) Immunofluorescence of U2OS repositioning cell lines in control (fed) condition, starved or starved in 5 nM A/C, fixed and stained for LC3 (green) and nuclei (blue). Confocal microscope, scale bar: 15 μm. (B) Quantification of co-localization of LC3 and LT1-mCherry-FRB* in starved U2OS and RPE-1 repositioning cell lines, showing fraction of LC3 overlapping with mCherry (autolysosomes). N=3 independent experiments. Ordinary one-way ANOVA was used. Lines: Mean +/- SD. (C) Immunofluorescence of U2OS repositioning cell lines, starved and then restimulated with or without 5 nM A/C for 30 min. Cells were fixed and stained for LC3 (green) and nuclei (blue). Confocal microscope, scale bar: 15 μm. (D) Total area (μm^2^) and number of LC3 puncta per cell in U2OS and RPE-1 repositioning cell lines. Cells were treated as in (C) and then fixed, imaged on an epifluorescence microscope and LC3 puncta quantified. N=3 independent experiments. Kruskal-Wallis test was used. (E) Western blot analysis of starved and re-fed U2OS cells. Cells were induced with 5 nM A/C (2h), starved (2h) and then restimulated (30 min). N=4 independent experiments. (F) Densitometric quantification of LC3-II. LC3-II and actin bands were quantified and their ratios were normalized to each starved sample. N=4 independent replicates, colors of data points indicate the replicates. Ordinary one-way ANOVA was used. Lines: Mean +/- SD.

To remove the potential noise of random spatial artifacts, we developed a dynamic clearance assay: cells were starved to induce autophagosome formation and subsequently re-fed with complete medium for 30 minutes to stimulate the autophagosomal clearance. Because of the slow and variable clearance kinetics observed in RPE-1 cells, we prioritized the U2OS line for subsequent characterization of autophagic flux. The U2OS line provided a more robust and quantifiable response. The assay revealed a significant accumulation of uncleared LC3 puncta ‒ measured by both count and total area ‒ in re-fed U2OS cells with peripheral lysosomes (**Figure 3 c–d**). This was confirmed biochemically by quantifying LC3-II, which corresponds to the lipid conjugated LC3 (**Figure 3 e–f**). Interestingly, a similar, albeit less pronounced, accumulation was observed microscopically in cells with perinuclear-targeted lysosomes but could not be confirmed by Western blotting. These data suggested that lysosomal clustering acted as a “trafficking bottleneck” for autophagy. Sequestering lysosomes to the cell periphery caused the cell to lose the spatial coverage necessary for efficient autophagosome-lysosome fusion, hindering global autophagic clearance.

We have explored several fundamental features distinguishing perinuclear and peripheral lysosomes in terms of their functional properties and specialization to distinct tasks within the cell. These roles must be mediated by dedicated lysosomal proteins, and our observations open the possibility of distinct proteomic signatures of each lysosomal population.

### Distinct proteomic profiles of perinuclear and peripheral lysosomes

The functional specializations we observed, specifically the spatial difference in pH, proteolytic activity, and exocytic competence, suggest that perinuclear and peripheral lysosomes have different molecular identities. To characterize these differences, we devised a proteomic strategy that uses our repositioning machinery as an enrichment tool. Unlike traditional LysoIP methods, which require an additional bait protein such as TMEM192 (Abu-Remaileh et al., 2017), our strategy utilizes the N-terminal 3×HA tags that are already present on our BicD2 and KIF5C motor constructs.

This approach enables the selective enrichment of lysosomes that are recruited by motor proteins upon A/C induction (**Figure 4 a**). Non-induced parallel samples served as stringent negative controls to account for nonspecific bead binding. Western blot analysis confirmed the efficient and specific enrichment of core lysosomal markers, such as LAMP2, TMEM192, BORCS6, and Rab7. Markers for other organelles were either undetectable or present at background levels (**Figure S3 b**). We observed that 3×HA-KIF5C was more effective than 3×HA-BicD2 at immunoprecipitating lysosomes, likely due to higher expression levels or reduced steric hindrance. While this difference was manageable in RPE-1 cells through careful normalization, it was more pronounced in U2OS cells. Thus, we focused our unbiased proteomic mapping exclusively on the RPE-1 line.

**Figure 4:**
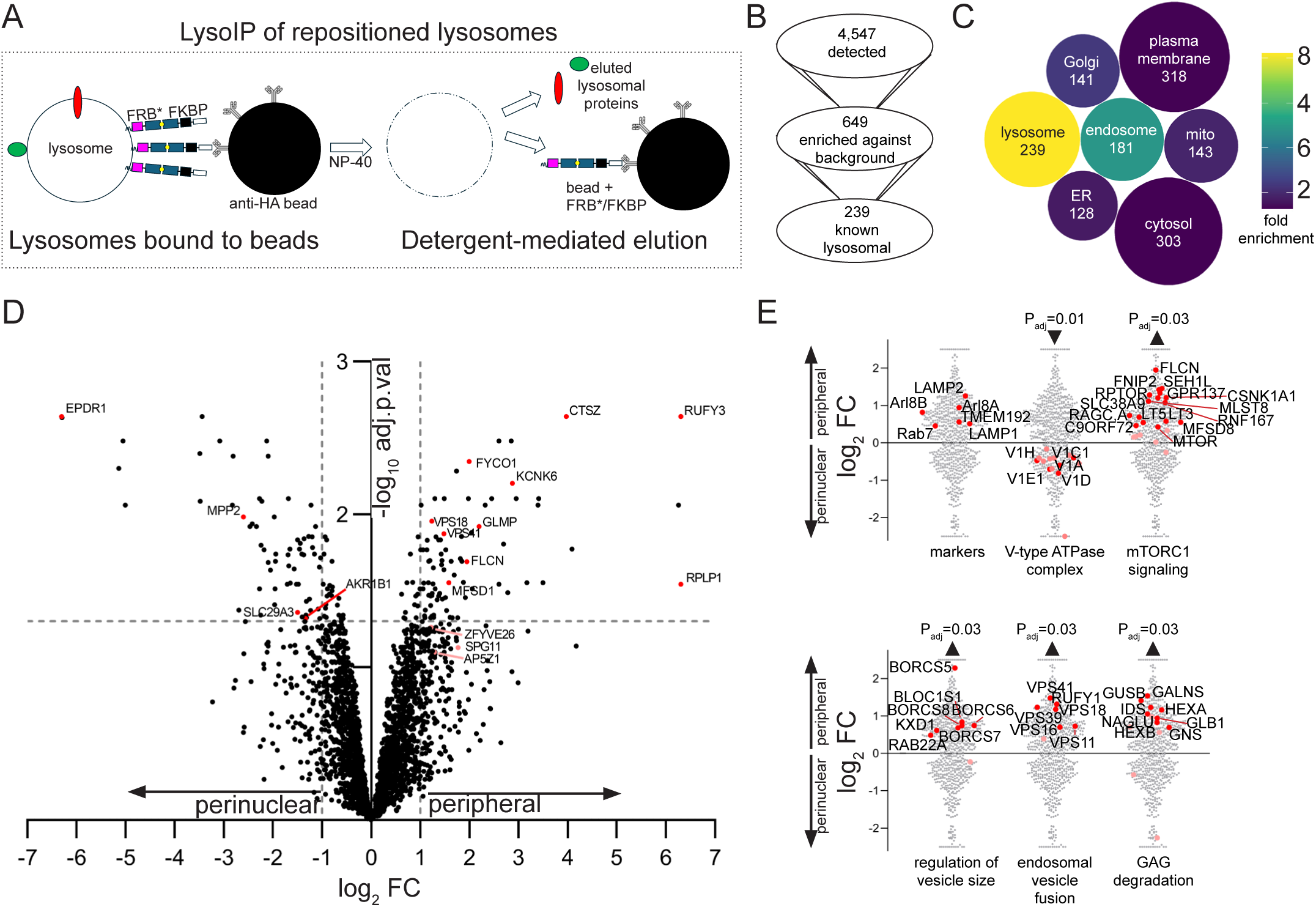
Proteomic characterization of perinuclear and peripheral lysosomes. (A) Schematic diagram depicting the LysoIP experimental setup for repositioning cell lines. After 4 h of A/C induction, endolysosomes were enriched from membrane fractions with anti-HA beads and endolysosomal proteins were then eluted in a detergent-containing buffer. (B) Overview of proteins detected by mass spectrometry (top), a subset of proteins enriched (P_adj_<0.05) against background control (middle) and a subset of proteins known to be lysosomal (bottom). (C) Proteins enriched over the background control sorted by organelle based on the COMPARTMENTS dataset. Color indicates fold-enrichment over a random scenario, size corresponds to the number of hits for each compartment. (D) Volcano plots of identified proteins, showing distribution of hits with their log_2_ fold change (FC) and -log_10_ adjusted P-value (calculated with DEqMS). Horizontal line at P_adj_=0.05, vertical lines at log_2_FC=1. Red: significant hits. Pink: additional hits explored in the text. (E) Enriched gene sets with corresponding adjusted P-value. Selected enriched gene sets from fast gene enrichment analysis (FGSEA) were visualized on scatter plots. Red points denote enrichment of gene set members forming the leading edge of FGSEA enrichment results. Pink: Other members of the same gene set. The “markers” plot is a selection of canonical lysosomal markers showing distinct enrichment patterns, specifically in LAMP2. The rest of the gene sets can be found using these references: GO:0033176 (proton-transporting V-type ATPase complex), GO:0038202 (TORC1 signaling) GO:0097494 (regulation of vesicle size), GO:0034058 (endosomal vesicle fusion), map00531 (glycosaminoglycan degradation).

To generate a comprehensive proteomic map, we performed label-free mass spectrometry on three independent biological replicates, each containing A/C-induced samples (BicD2 and KIF5C) and their respective non-induced controls. We showed that the BicD2 and KIF5C triplicates clustered together, indicating experimental reproducibility (**Figure S3 c**). We then employed the DEqMS algorithm developed specifically for differential expression analysis, which accounts for the number of peptides used for the quantification of each protein in label-free proteomics to calculate adjusted P-values (Zhu et al., 2020). The DEqMS variance output was used for the BicD2 / KIF5C and the BicD2 / control comparisons, but the mean-variance trend did not meet the assumptions required for the KIF5C / control comparison, thus the standard limma output was applied there (**Figure S3 d**).

Out of 4,547 detected proteins, 649 were significantly enriched in either induced BicD2 or KIF5C samples compared to the controls (**Table 2**). Although non-specific protein recovery is an inherent challenge in LysoIP protocols (Akter et al., 2023), including non-induced controls allowed us to subtract these background contaminants computationally. We identified specific organelle enrichment using these proteins as compared to the COMPARTMENTS dataset (Binder et al., 2014) and revealed robust enrichment of canonical lysosomal and endosomal markers. Specifically, the identification of 239 known lysosomal proteins constitutes about 27% of the annotated human lysosomal proteome. Conversely, markers for other organelles were reduced to baseline levels, confirming the high subcellular specificity of our spatial enrichment strategy (**Figure 4 b–c**).

The comparison of perinuclear and peripheral lysosomes revealed 89 proteins significantly elevated in the perinuclear sample and 108 proteins in the peripheral lysosomes (**Figure 4 d**). In addition to these individual proteins, an unbiased analytical approach can track global changes by measuring enrichment of functional subsets. We used gene-set enrichment (FGSEA) to identify differentially regulated pathways within these lysosomes (**Table 2**). Among these pathways, the V-ATPase complex was significantly associated with the perinuclear lysosomes, serving as an important quality control and offering a direct explanation for the increased acidity of this lysosomal pool. The peripheral lysosomes are enriched in pathways controlling lysosomal dynamics, such as the regulation of lysosomal size (de Araujo et al., 2020) represented primarily by BORC complex components (Filipek et al., 2017; Yordanov et al., 2019; de Araujo et al., 2026) or fusion (i.e. HOPS complex). Additionally, glycosaminoglycan degradation pathways are also further represented in the periphery, and their signaling function is underpinned by the enrichment of the mTORC1 signaling components (**Figure 4 e**).

To identify high-confidence candidates for spatial specialization, we isolated 87 proteins that exhibited significant differential enrichment between locations while maintaining a clear signal over the non-induced background. This filtering process revealed distinct molecular profiles for each subpopulation. The perinuclear region was characterized by the enrichment of EPDR1, a luminal protein of unknown function (Wei et al., 2019), as well as the nucleoside/urate transporter SLC29A3/ENT3 (Baldwin et al., 2005; Matake et al., 2025). In comparison, peripheral lysosomes were enriched in the GLMP/MFSD1 dipeptide transporter complex (Jungnickel et al., 2024) as well as specialized cathepsins, including CTSZ and CTSH. Moreover, some proteins fell below the threshold of significance, but their enrichment was obvious when considering their mutual physical association; these include the peripheral-enriched SPG11/SPG15/AP5 complex involved in lysosomal tubulation (pink data points, **Figure 4 d**). Established pan-lysosomal markers lacked significant enrichment toward neither the periphery nor the perinuclear group except for LAMP2 which showed a significant affiliation with the peripheral lysosomal population (**Figure 4 e**, top left).

There was also significant enrichment of endoplasmic reticulum (ER)- and Golgi-associated proteins in the perinuclear sample, likely a consequence of an increased amount of contact sites with these organelles (Barral et al., 2022). Fold-change ratios of proteins assigned to the ER or the Golgi apparatus are skewed towards the perinuclear lysosomal sample (**Figure S3 e**) and GO terms associated with the ER/Golgi are over-represented in the perinuclear lysosomes (**Figure S3 f**). Indeed, ER network is particularly dense in the perinuclear region. However, densities of ER tubules and sheets can be also induced by repositioning the lysosomes to the periphery (**Figure S3 g**), demonstrating the known role of the lysosomes in the organization of the ER network (Lu et al., 2020).

### Protein localization in wild-type cells follows trends from repositioning cell lines

Finally, we tested whether proteomic results from the repositioning cell lines can be extrapolated to wild-type cells. We used immunofluorescence and co-stained selected hits together with LAMP1, a marker which almost equally localizes to the perinuclear and peripheral lysosomes. We then used the LAMP1 channel to create a lysosomal mask and performed shell analysis with concentric rings around the nucleus. In the end, 12 equally large regions of increasing distance from the nucleus were calculated and the signal intensity plotted on histograms (**Figure S4 a‒d**) or relative to LAMP1 (**Figure 5**). V1B2, a member of the V-ATPase gene set enriched in the perinuclear lysosomes, served as our positive control. We observed an enrichment of V1B2 signal in the perinuclear lysosomes relative to LAMP1 in U2OS and RPE-1 cells (**Figure 5 a**). We also plotted LAMP2 against LAMP1 and found that it was indeed enriched on peripheral lysosomes in RPE-1 cells; however, this was not the case in U2OS cells, where LAMP1 and LAMP2 profiles did not deviate (**Figure 5 b**).

**Figure 5:**
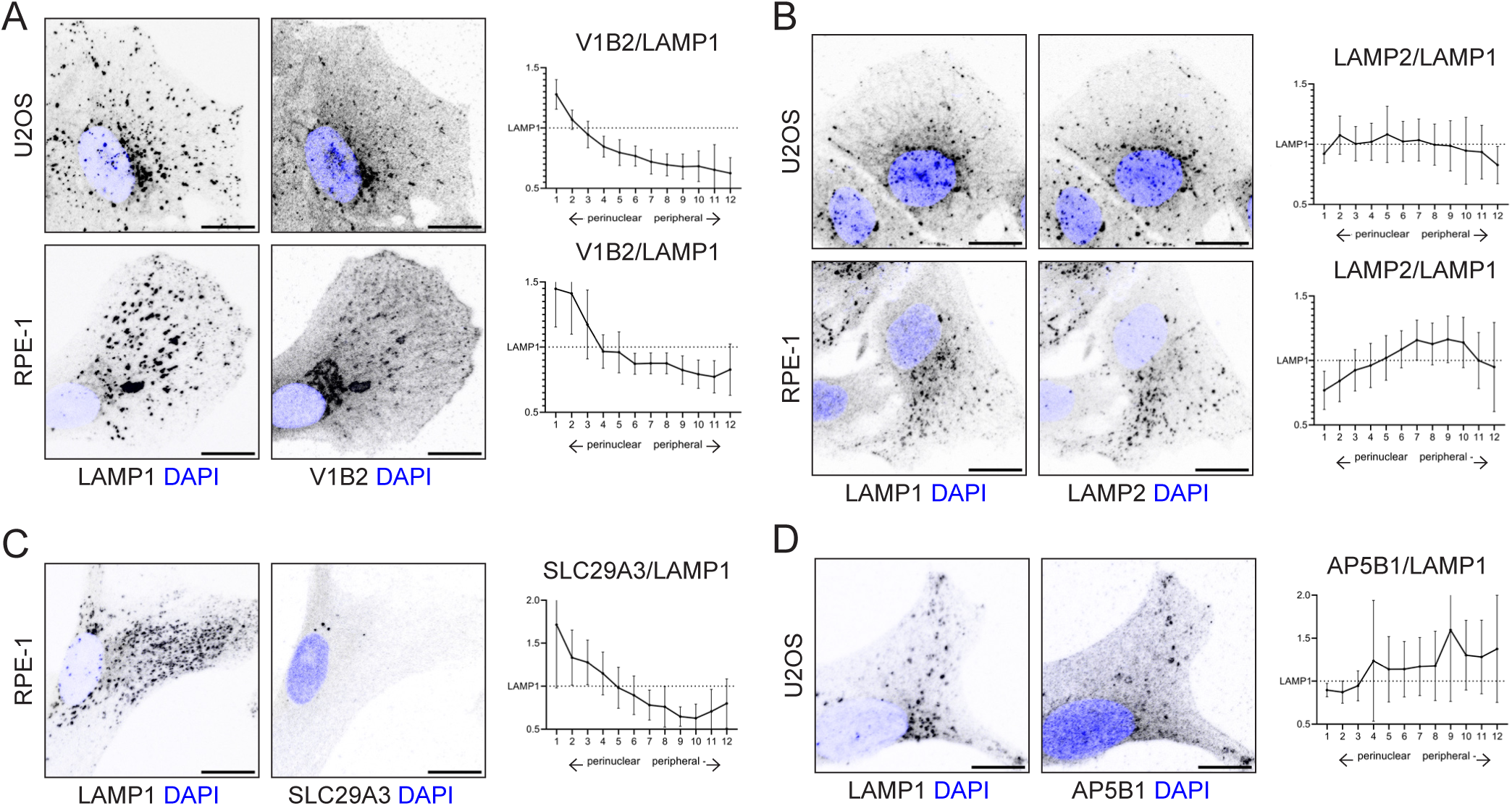
Immunofluorescence staining of perinuclear and peripheral lysosomal proteins. (A, B, C, D) Left: Immunofluorescence of U2OS and RPE-1 WT cells using antibodies against LAMP1 and one of the validated hits (V1B2, LAMP2, SLC29A3, or AP5B1). Greyscale inverted images overlaid with nuclear stain (blue, DAPI). Confocal microscope, scale bars: 15 μm. Right: Analysis of antibody signal distribution from the perinuclear to the peripheral areas of the cell. Cells were divided into 12 equally sized areas of increasing distance from the nucleus (1=most perinuclear, 12=most peripheral) and % of total signal in each area was normalized to % signal of LAMP1 in that area of the cell.

We then strived to use this technique to characterize lesser-known lysosomal proteins identified in the proteomic data. Although large-scale screening was limited by the availability of antibodies and high cytosolic backgrounds, as seen with HOPS components, we successfully identified several key markers. For the perinuclear pool, the SLC29A3 protein was found almost exclusively in a specific population of juxtanuclear lysosomes in RPE-1 cells (**Figure 5 c**). In U2OS cells, the SLC29A3 signal appeared diffuse, likely due to lower endogenous expression levels. To confirm peripheral enrichment, we targeted the SPG11/SPG15/AP5 complex. Although robust antibodies for the primary proteomic hits were lacking, we detected the AP5B1 subunit within the peripheral lysosomes of wild-type U2OS cells (**Figure 5 d**), which provides indirect validation of this complex. Furthermore, cathepsin Z (CTSZ) was found to be preferentially localized to dense patches at the plasma membrane of RPE-1 cells (**Figure S4 e**), which is consistent with the specialized exocytic capacity of the peripheral pool. However, other candidates, such as EPDR1 and GLMP, could not be validated due to the absence of antibodies suitable for immunofluorescence analyses.

These findings show that the proteomic signatures identified in our repositioning lines faithfully predicted the special distribution of lysosomal proteins in wild-type cells. Because these spatial patterns, such as the perinuclear enrichment of V_1_ V-ATPase subunits and the peripheral localization of CTSZ, persist in cells lacking exogenous motor recruitment, we conclude that our findings reflect authentic lysosomal specialization rather than artifacts of motor activity. The challenges we faced with antibody-based validation highlight the need for unbiased mass spectrometry to understand the true molecular landscape of lysosomal heterogeneity, which traditional imaging methods often miss.

## Discussion

Despite substantial progress in our understanding of lysosomal repositioning, it remains unclear how spatially distinct lysosomal pools differ in their molecular composition and whether repositioning alone is sufficient to drive functional adaptation. In particular, the lack of tools to selectively manipulate lysosomal localization without broadly perturbing cellular physiology has limited our ability to disentangle the direct consequences of lysosomal positioning from secondary stress or starvation responses. In this study, we characterize functional consequences of repositioning lysosomes to the perinuclear or peripheral area and provide the first direct proteomic comparison of perinuclear and peripheral lysosomes.

Our repositioning system enables inducible recruitment of motor proteins to lysosomes within minutes, their efficient repositioning after 30 minutes, and functional consequences such as pH modulation observed within 4 h. The fact that lysosomal pH changes after lysosomal repositioning so rapidly suggests that endolysosomes acutely respond to, and adapt to, their new cellular environment within this time frame. Long-term changes in gene expression, e.g. mediated by a CLEAR response, are unlikely to have a strong effect at a 4-h time point. While the “master regulator” of the CLEAR network, TFEB (Transcription Factor EB), can translocate to the nucleus rapidly in response to stress (e.g., within minutes of starvation or oxidative stress), the subsequent transcription and accumulation of target lysosomal and autophagic genes often follow a slower kinetic profile (Sardiello, 2016; Li et al., 2021). Thus, our system provides a powerful tool to dissect the direct effects of lysosomes residing in perinuclear or peripheral locations, as well as the immediate functional consequences of their enrichment at these sites.

In RPE-1 cells, the elevated pH of peripheral lysosomes is associated with reduced V_1_-ATPase presence and significantly lower proteolytic activity. The specific adaptive mechanisms that promote V_1_–V_O_ holoenzyme assembly when lysosomes relocate to the perinuclear region are unclear. Although the mRAVE complex facilitates this assembly in MEFs and neurons (Ratto et al., 2022; Verma et al., 2025), its components were not enriched in our lysosomal proteome. This discrepancy suggests the existence of alternative pathways, perhaps cell-type specific, for spatial pH regulation. Taken together, these results imply that lysosomal adaptation, whether through the delivery of new machinery or the removal of existing proteins, is a highly regulated process that actively remodels the organelle to suit its local microenvironment.

Our results also provide strong indications that mTORC1 signaling is not significantly influenced by lysosomal position in our cell lines, and it is possible that the signaling outcomes observed in some studies (Korolchuk et al., 2011; Clippinger and Alwine, 2012; Walton et al., 2019; Dang et al., 2026) could rather be attributed to side effects of lysosomal repositioning strategies. Interestingly, peripheral lysosomes in RPE-1 cells are enriched with specific components of the mTORC1 regulatory machinery, such as the amino acid sensor SLC38A9 and the GEF folliculin. This spatial partitioning suggests that, although peripheral lysosomes have reduced degradative capacity, they may be specialized for nutrient sensing and signal transduction. This specialization could be particularly relevant during prolonged physiological shifts. For instance, mild starvation exceeding 4 h has been shown to modulate the metabolic impact of lysosomal distribution (Dang *et al*., 2026.). Our inducible system provides a unique opportunity to test directly whether these spatial signatures are primary drivers of the starvation response.

Beyond signaling, our study provides clear evidence that lysosomal positioning is a critical determinant of autophagic flux. We found that sequestering lysosomes in the periphery leads to a significant decrease in autolysosome formation during starvation and greatly disrupts autophagosome clearance upon nutrient restimulation. These findings suggest that spatial segregation of lysosomes creates a “diffusion-limited” environment that reduces the probability of encounter between lysosomes and autophagosomes. This conclusion aligns with previous stochastic modeling and experimental studies (Cardoso et al., 2009; Laopanupong et al., 2021). We could microscopically detect the same outcome in cells with the lysosomes repositioned to the perinuclear area, and indeed this phenomenon was also previously reported (Jia et al., 2017). In our hands, this was less robust than with the peripheral lysosomes and not observed biochemically. This indicates that most autophagosome-lysosome fusion events naturally occur within the perinuclear space. Thus, this region is less sensitive to additional lysosomal crowding than the sparsely populated periphery.

Finally, we also explored lysosomal exocytosis and observed an increase in RPE-1 cells with lysosomes repositioned to the periphery. Again, this could be attributed to the higher probability of encounters, in this case between the lysosomes and the plasma membrane. Regulating exocytosis may have important functional consequences for the cell’s ability to degrade the extracellular matrix and migrate. While U2OS and RPE-1 cells are impractical models for cell migration due to their rather sedentary nature, these phenomena were recently studied in depth in melanoma cells using a system similar to ours (Jerabkova-Roda et al., 2025).

One definitive strength of our methodology is our ability to isolate repositioned lysosomes and compare their proteomic landscapes directly and without bias. Our data show that perinuclear lysosomes are configured to recycle metabolites; in addition to having a high density of V_1_ V-ATPase subunits, they contain the nucleoside transporter SLC29A3 (ENT3). These findings suggest that the perinuclear pool serves as a specialized hub for the breakdown and subsequent efflux of nucleic acid components. Additionally, the increased abundance of Golgi and ER proteins in this fraction likely reflects the dense network of organelle contact sites or elevated retrograde trafficking characteristic of the juxtanuclear environment. In contrast, the peripheral proteome is specialized for extracellular interaction. We observed significant enrichment of machinery dedicated to degrading the extracellular matrix, specifically glycosaminoglycans, alongside specialized secretory proteases, such as cathepsin Z (CTSZ). This proteomic signature directly supports the role of peripheral lysosomes as exocytic vesicles. Finally, our independent validation of the SPG11/SPG15/AP5 complex on peripheral lysosomes identifies a spatially restricted niche for this machinery. This provides a molecular basis for its specialized role in membrane trafficking and organelle homeostasis. This complex associates with curved membranes and promotes lysosomal reformation (Mai et al., 2025), which correlates with a tubular, dynamic nature of the peripheral lysosomes (Rayens et al., 2022). Surprisingly, LAMP2, a frequently used marker for lysosomes, is also enriched in the peripheral lysosomes in the RPE-1 cells. The spatial separation of LAMP1 and LAMP2 is even more pronounced in human neurons, where LAMP2 shows limited overlap with LAMP1 and vesicles rich in LAMP2 are more dynamic than those primarily coated with LAMP1 (Grochowska et al., 2023; Abouward et al., 2026). This necessitates caution when using LAMP2 as a lysosomal marker in certain cell lines.

Our data did not provide conclusive evidence supporting the hypothesis that Rab7A and Arl8b are strictly segregated between perinuclear and peripheral lysosome populations (Johnson et al., 2016; Schleinitz et al., 2023). Although both GTPases were identified as high-abundance components in our RPE-1 proteome, neither was significantly enriched in one spatial pool over the other. Independent validation by immunofluorescence remains technically challenging because of the substantial Rab7A cytosolic fraction and the lack of reliable commercial Arl8b antibodies. Additionally, while recent studies have identified distinct phosphoinositide signatures, specifically varying PI(3)P-to-PI(4)P ratios between spatial pools (Ebner et al., 2023), our current proteomic methodology was not used here to detect these lipidomic differences. Future studies integrating spatial lipidomics with our protein enrichment system are necessary to fully characterize these organelle niches.

Although our inducible systems are a significant improvement over traditional methods for studying lysosomal positioning, we recognize the limitations of engineered cellular models. Although the LT1 tag is designed to minimize structural disruption associated with full-length LAMP1 overexpression, it remains an exogenous modification that could theoretically influence organelle behavior. Similarly, recruiting KIF5C and BicD2 domains introduces the potential for unintended interactions with endogenous motor regulators. However, the indistinguishable proliferation rates and high viability of our repositioned lines imply that global microtubule-based traffic remains largely unperturbed. Additionally, the substantial time difference between rapid transport (30 minutes) and functional adaptation (4 hours) indicates that the observed changes in pH and proteolytic activity are a result of the new intracellular environment rather than the transport process itself. Although we used a rapalog with minimal inhibitory effects on mTORC1, the need for chemical induction requires careful experimental control. Additionally, while acute, forced repositioning is a powerful strategy for isolating spatial variables, it is, by nature, an induced state. Our data demonstrate that lysosomes adapt functionally and proteomically to their new niches; however, certain “positional memories” from their original locations may persist. Lastly, we utilized LAMP1 for normalization based on its equal distribution in our proteomic datasets. However, we acknowledge that a subset of LAMP1 may localize to non-lysosomal compartments in specific contexts (Cheng et al., 2018). Nonetheless, by integrating these considerations into our analysis, we provide a robust framework for dissecting the spatial logic of the lysosome.

In summary, although we successfully validated key candidates using immunofluorescence, the scarcity of high-affinity antibodies highlights why unbiased mass spectrometry is the preferred method for resolving lysosomal heterogeneity. Our study provides the first proteomic characterization of lysosomes with spatial resolution by bypassing the inherent limitations of traditional imaging. These datasets offer the scientific community a valuable resource, providing a molecular roadmap to investigate how positioning dictates organelle function. In addition to these findings, we are making our unique collection of engineered plasmids and cell lines fully available to the scientific community. We anticipate that adopting this inducible system in diverse, strongly polarized cells (e.g., neurons) or highly motile cells will be instrumental in deciphering how spatial logic adapts to meet complex physiological demands. Ultimately, by providing the fundamental data and tools for further expansion, this work establishes a new framework for investigating how the geography of the cell governs its function in health and disease.

## Materials and methods

### Antibodies and primers

Antibodies and primers used in this study are listed in **Table 1**.

### Plasmids and molecular cloning

The LT1-mCherry-FRB* vector was engineered from LAMP1-mCherry-FRB* (T2098L), which was a gift from Anthony Galione (Addgene plasmid # 207145) (Davis et al., 2020). LT1 tag was introduced in exchange for LAMP1 via an oligo encoding the 39 N-terminal amino acids of human LAMTOR1 flanked by EcoRI and SalI+linker sites. For generation of stable cell lines, retroviral vectors pMXs-Puro and pMXs-EF1-Blasti (Kitamura et al., 2003) were a gift from Vladimír Kořínek and pMXs-IRES-Hygro was a gift from Richard Possemato (Addgene plasmid # 164142). Vectors pMXs-EF1-Zeo and pMXs-EF1-G418 were made by PCR-amplifying Zeocin and G418 resistance genes and inserting them into the selection cassette of pMXs-EF1-Blasti. The open reading frame of LT1-mCherry-FRB* was PCR-amplified and inserted via BamHI and NotI sites into pMXs-IRES-Hygro and pMXs-EF1-G418. For the FKBP part, we started with pBa-tdTomato-flag-BicD2 594-FKBP encoding mouse Bicaudal D2 1-594aa, which was a gift from Gary Banker and Marvin Bentley (Addgene plasmid # 64205) (Bentley et al., 2015). First, a 3×HA tag was added between BsrGI and EcoRV sites by PCR amplification of the sequence from pcDNA3-3HA-TurboID (Addgene plasmid # 107171) (Branon et al., 2018). Then, 3×HA-BicD2-FKBP was PCR-amplified and subcloned into pMXs-EF1-Zeo and pMXs-EF1-Blasti via BamHI and NotI sites. This vector was used to create a corresponding KIF5C counterpart. First, an EcoRI site was engineered in a linker upstream of BicD2 with a two-step PCR reaction using two “external” primers to delineate a larger amplified section and two incompletely homologous “internal” primers creating the EcoRI site. Finally, BicD2 was replaced with rat KIF5C motor domain 1-559aa by PCR amplification from pBa-KIF5C 559-tdTomato-FKBP (Addgene plasmid # 64211) with EcoRI and RsrII sites.

Lyso-TORCAR was a gift from Jin Zhang (Addgene plasmid # 64929) (Zhou et al., 2015). For retroviral expression, the open reading frame was subcloned via NotI sites into pMXs-EF1-Blasti. All vectors were verified by Sanger sequencing.

### Cell culture and treatments

All cell lines were maintained at 37 °C with 4% CO₂ and 98% humidity, and their culture media were regularly tested for mycoplasma contamination. U2OS and RPE-1 cells were grown in high glucose DMEM with GlutaMAX (Thermo Fisher) supplemented with 10% fetal bovine serum FBS Superior (Sigma Aldrich) and 1% Pen Strep solution (Merck) (“complete medium”). Growth medium for MEFs was identical except it contained 20% FBS Superior. All cells were routinely passaged with a Trypsin-EDTA solution in PBS (Sigma Aldrich) and discarded within 10 passage cycles; for long-term storage, cells were frozen in FBS with 10% DMSO and stored in -80 °C for up to 3 months or in liquid nitrogen.

For lysosomal repositioning, A/C heterodimerizer (AP21967, Takara Bio), prediluted in ethanol to a 1000× or 10000× stock, was added to cell culture medium at 5 nM final concentration. Equal volumes of ethanol were used for a control treatment.

For starvation, cells were washed twice with PBS and incubated for 2h in Dulbecco’s MEM Ham’s F12 with HEPES, NaHCO_3_, without amino acids (USBiological), pH 7.4, without addition of serum (this will be referred to as “DMEM AA-FBS-“).

### Generation of stable cell lines

Retroviral vectors encoding the repositioning constructs were transfected into the Phoenix packaging cell line using Lipofectamine LTX with Plus reagent (Thermo Fisher) in full medium. Medium was exchanged 4 h after transfection and supernatants containing virus particles were collected 48 h after transfection. Transduction was done on freshly seeded cells at approximately 50% confluence, by incubating cells with the supernatant supplemented with 8 μg/mL Polybrene (Sigma Aldrich) for 24 h. Transduced cells were then selected with fresh medium supplemented with appropriate selection antibiotics. The latter varied because wt RPE-1 cells are already resistant to hygromycin and puromycin and MEF cells were incompletely eliminated with zeocin even after 2 weeks of cultivation. Therefore, cells were first transduced with constructs expressing LT1-mCherry-FRB* (U2OS: hygromycin resistance, 400 μg/mL hygromycin; RPE-1: G418 resistance, 200 μg/mL G418; MEF: hygromycin resistance, 200 μg/mL hygromycin). After negative control was eliminated by the antibiotic, the processed was deemed complete. Cells were then transduced with constructs expressing 3×HA-BicD2-FKBP or 3×HA-KIF5C-FKBP (U2OS and RPE-1: zeocin resistance, 300 μg/mL; MEF: blasticidin resistance, 10 μg/mL blasticidin) and again selected in parallel to a negative control. The resulting batch cell lines showed heterogeneous expression levels of both constructs and varying ability to reposition. Therefore, single cell clones, which showed adequate expression of both constructs and a robust lysosomal response to A/C treatment, were selected for further experiments.

### Microscopy

Screening of clones (**Figure 1 c**) and high-throughput microscopy for LC3 puncta quantification (**Figure 3 d**) was done on a Zeiss AX10 epifluorescence microscope with a 40× Plan-Neofluar objective (NA 1.3, oil immersion) and a Prime BSI Express sCMOS camera. In all other figures, confocal microscope LSM980 (Zeiss) with the LD LCI Plan-Apochromat 63×/1.2 Imm Corr DIC M27 objective (glycerol immersion) was used. Appropriate lasers (405 nm for DAPI/Hoechst, 488 nm for EGFP/Alexa 488, 561 nm for mCherry/Alexa 568, 639 nm for Alexa 647) were organized in separate tracks to allow perfect separation of channels; an additional brightfield channel was captured where appropriate (T-PMT). Scanning was done with 1.0× Nyquist sampling at a pixel time of 0.77 μs and Z-Stacks were captured with optimal slicing. 8-bit. CZI images were saved and processed in ImageJ 1.54r.

Airyscan setup (SR-4Y) was employed for high-resolution microscopy (**Figure 1 d**) and these images were then deconvolved (CMLE) in Huygens Professional 25.10. Live imaging was done in a humidified atmosphere (95 %) with 4 % CO_2_ and at 37 °C. In this case, microscope settings were optimized for quick acquisition speed (see below for each method using live cell imaging).

IncuCyte Live Cell Analysis System was used for cell growth analysis (growth curves) and for **Figure S1 e‒f**. Cells were seeded onto 24-well dishes 24 h before the start of imaging. Once complete, “basic analysis” mode was used for quantification of area covered by cells over time as a proxy of cell proliferation speed.

### Shell analysis

For **Figure 1 e**, BicD2 and KIF5C cell lines were seeded 24 h before imaging at 20,000/well in an Ibidi 8 Well µ-Slide. Before induction with A/C, growth medium was replaced for Fluorobrite DMEM (Thermo Fisher) supplemented with 2 mM L-glutamine (Thermo Fisher), 10% FBS and 1 µg/mL Hoechst 33258. The slide was equilibrated on the microscope stage. Z-stacks of confocal images were then captured at 5 min intervals. For quantitative analysis, maximum intensity projections were made in ImageJ, crosstalk of Hoechst 33258 into the mCherry subtracted, and cell areas were manually saved as regions of interest (ROIs). The mask of the nucleus in each ROI was then automatically detected by applying median filter (radius 20) and default thresholding. We then used an iterative shell enlargement around the nucleus in accordance with earlier methodological approaches (Starling et al., 2016). For perinuclear area, band around the nucleus was incrementally enlarged until it equaled 20% of the total cytosolic area, and then integrated density of mCherry signal was measured there. For the peripheral area, the integrated density of mCherry inside the farthest 20% of cytosol from the nucleus was quantified. Both values were then expressed as % of total cellular mCherry signal.

A similar but distinct workflow was employed for the quantification of antibody signal in final hit validation staining. In this case, representative images of fixed and immunofluorescence-stained coverslips with WT U2OS or RPE-1 cells were acquired in Z-Stacks, merged with maximum projection, and cellular ROIs were manually selected in ImageJ. An automated script was then used to threshold lysosomes using LAMP1 staining (with the “Moments” automatic threshold to reduce bias). For the rabbit LAMP1 antibody, background was automatically subtracted from this channel (50 px radius) before thresholding. Subsequently, integrated density of LAMP1 and a given validated hit was measured inside the lysosomal ROI by incrementally expanding the nuclear ROI by 1 pixel until the whole area of the cell was included in this expanding circle. For each measured cell, this workflow created one table with an incrementally expanding cytosolic area and a corresponding integrated density measurement. These were finally processed in R by creating 12 cytosolic regions with identical area as a basis for the histogram plotting of intensity (script writing in R was aided by Gemini Advanced).

### Immunofluorescence

For immunofluorescence, cells were seeded onto coverslips 24–48 h in advance to reach approximately 25% confluence on the day of the experiment. Cells were fixed in 4% paraformaldehyde (PFA) in CB buffer (10 mM PIPES pH 6.8, 150 mM NaCl, 5 mM EGTA, 5 mM glucose, 5 mM MgCl_2_) and washed 2× with CB and 1× with 50 mM ammonium chloride in CB. A 30-min blocking step was then performed with 3% bovine serum albumin and 0.025% saponin in CB. Coverslips were then serially incubated with primary antibodies diluted in blocking buffer, washed, and incubated with fluorescent-conjugated secondary antibodies in blocking buffer. Finally, coverslips were mounted in Mowiol with DAPI.

For V1B2/LAMP1 simultaneous staining, cells were first stained with anti-LAMP1 and a corresponding secondary antibody as indicated above. Then, cells were refixed with PFA, treated with 0.025% SDS in PBS for 5 min, washed thoroughly, and incubated overnight with the anti-V1B2 antibody. As a last step, cells were incubated with the corresponding secondary antibody. SLC29A3/LAMP1, AP5B1/LAMP1 or CTSZ/LAMP1 combinations were processed similarly but using 0.075% SDS (SLC29A3, CTSZ) or 0.5% SDS (AP5B1), respectively.

### Lysosome pH measurement

BicD2 and KIF5C cell lines were seeded 48 h before imaging at 10,000/well in an Ibidi 8 Well µ-Slide. Next day, LysoSensor Yellow/Blue Dextran 10,000 MW (Thermo Fisher) was added to all wells at 1 mg/mL. Cells were washed and induced 4 h before the experiment with 5 nM A/C in FluoroBrite DMEM (Thermo Fisher) supplemented with 10% FBS and 2 mM L-glutamine (Thermo Fisher). Cells were then equilibrated on a preheated confocal microscope, and lysosomal patches were captured in two channels enabling ratiometric measurement (ex 405 nm/em 411–499 nm or 508–587 nm). For each experiment, a pH calibration curve was also established to convert fluorescence ratios to pH (Johnson et al., 2016) by serial incubation of cells in K^+^ isotonic buffer (143 mM KCl, 5 mM glucose, 1 mM MgCl_2_, 1 mM CaCl_2_, 20 mM HEPES) of increasing pH (4.5–7.5 at 0.5 increments), supplemented with 10 μM nigericin (Sigma-Aldrich) and 5 μM monensin (Sigma-Aldrich).

### Magic Green assay

BicD2 and KIF5C cell lines were seeded 48 h before imaging at 10,000/well in an Ibidi 8 Well µ-Slide and supplemented with 0.2 mg/mL Dextran-Alexa 647 (Thermo Fisher) approximately 28 h before the experiment. Cells were then washed twice with PBS and incubated with fresh complete medium and A/C 4 h before the experiment. Control samples were incubated with 200 nM bafilomycin for 2 h. Magic Green reagent (Green Cathepsin B Kit, Bio-Rad) was added at t=0, at a recommended concentration (250× from the DMSO stock) resuspended in DMEM + 0.5% FBS (15 min, 37 °C), and then replaced with fresh complete medium. Cells were again incubated for 15 min in a live imaging chamber and then immediately imaged. Images were processed in ImageJ. Integrated density of Magic Green in lysosomal patches was first normalized to the integrated density of Dextran-Alexa 647 and then normalized to the median of each BicD2 replicate.

### DQ Ovalbumin assay

DQ Ovalbumin plate reader assay was adapted from a previously published protocol (Iwai et al., 2025). Repositioning cell lines were seeded in 6-well dishes at 2.5×10^5^/well (U2OS) or 2×10^5^/well (RPE-1) in a complete medium. A/C was added 4 h before the experiment. Bafilomycin (200 nM) for control treatments was added 1 h before the experiment. The next day, 20 μg/mL DQ (Green) Ovalbumin (Thermo Fisher) in 1 mL of DMEM supplemented with 0.5% FBS was added to each well and incubated for 4 h (with or without A/C). Dishes were then placed on ice, washed 2× with ice-cold PBS, and cells were lysed in 150 μL KPBS + 0.5% Triton X-100, pipetted thoroughly, and lysates were cleared by a 5-min centrifugation at 16,000 × g (4°C). 100 μL of cleared cell lysate was directly measured in a 96-well on a Spark microplate reader (Tecan, excitation: 485 nm with a bandwidth of 20 nm, emission: 525 nm with a bandwidth of 20 nm). Fluorescence readouts were then normalized to the corresponding non-induced control lysate (without A/C).

### Exocytosis assay

Repositioning cell lines were seeded in 12-well dishes at 0.5×10^5^/well (U2OS) or 0.4×10^5^/well (RPE-1) in a complete medium. After 24 h, they were washed 2× in PBS and incubated in 600 μL DMEM without serum for another 24 h (with or without A/C). Positive controls were made by treating cells with 2 μM ionomycin for 5 min. Supernatant was then collected and centrifuged at 600×g (5 min, 4°C) to remove floating cells (cleared supernatant). In the meantime, cells grown in 24-well plates were left shaking in 500 μL PBS + 0.2% Triton X-100 for 5 min in room temperature to release cellular enzymatic activity (cell lysate). Enzymatic reactions in 96-wells consisting of 50 μL 10 mM 4-Nitrophenyl N-acetyl-β-D-glucosaminide (Merck) in 0.1 M sodium citrate (pH 4.6, containing 0.2% BSA) and 25 μL cleared supernatant or 4 μL cell lysate were incubated for 24 h / 37°C. Reaction was then stopped with 200 μL stop solution (400 mM glycine pH 10.4) and absorbance of 4-Nitrophenol at 405 nm was measured on a Spark microplate reader (Tecan). For the ionomycin samples, absorbances of the supernatant were normalized to the absorbance of each untreated control supernatant. For repositioning experiments, absorbances of the supernatant were normalized to the absorbance of the corresponding cell lysate, and then to the mean absorbance of each uninduced control.

### Autophagy induction and quantification

Quantification of LC3/mCherry overlap (autolysosomes) was done by treating cells with A/C or control for 2 h and then starving them for 2 h in DMEM AA-FBS-with or without A/C (see cell culture for details). Autophagosome clearance in re-fed conditions was induced similarly, followed by a 30 min cultivation in complete medium with or without A/C. For the calculation of Manders coefficient, cell ROIs were manually saved in ImageJ. For each cell, LC3 signal and mCherry signal were thresholded using pre-trained classifiers in Labkit (Arzt et al., 2022) and Manders tM2 was calculated with JACoP (Bolte and Cordelières, 2006). Similarly, LC3 puncta in the re-fed conditions were quantified by thresholding the LC3 channel with a pre-trained Labkit classifier.

### Lyso-TORCAR measurements

Lyso-TORCAR FRET probe for mTORC1 activity was stably expressed in U2OS BicD2 and KIF5C and RPE-1 BicD2 and KIF5C cells via a retroviral vector encoding the construct. Polyclonal cell lines (to prevent clonal artifacts) were used. Cells were seeded 48 h before imaging at 10,000/well in an Ibidi 8 Well µ-Slide and induced with A/C 4h before the experiment. For a starved control, cells were washed with PBS twice and incubated with DMEM AA-FBS-supplemented with A/C for 2h before the experiment. mCerulean was excited with a 445nm laser and emission was detected simultaneously in the mCerulean window (464–499 nm) and in the YPet window (516–552 nm). FRET ratio was calculated by manually selecting perinuclear and peripheral patches as ROIs in ImageJ and measuring their integrated density in each channel. No normalization was introduced among replicates.

### Lysosome enrichment for MS (LysoIP)

RPE-1 BicD2 and KIF5C cell lines were each seeded in sixteen 15cm dishes in DMEM with 10% serum 48–72 h before harvest, to reach 0.7×10^7^ (RPE-1) or 1×10^7^ (U2OS) cells/plate at the day of the experiment. Half of the dishes were induced by addition of A/C and half were kept as controls. A/C was also included at all steps for the induced cells. Cells were washed once with 10 mL PBS and then carefully scraped into 5 mL PBS with a rubber policeman. Cell suspensions were then centrifuged once at 120 × g for 5 min, resuspended in 2 mL KPBS (136 mM KCl, 10 mM KH_2_PO4, pH 7.25, 10 μg/mL aprotinin, 1 μg/mL pepstatin, 10 μg/mL Leupeptin, 0.4 mM Pefablock SC), centrifuged again at 444 × g for 10 min, resuspended in 3–5× pellet volume KPBS and moved to a 15cm falcon tube. A post-nuclear supernatant (PNS) was then prepared by passing the cell suspension 30× through a 25-gauge needle pointing to the wall of the falcon tube and centrifugating at 1000 × g for 10 min. Concentration of PNS was determined by Bradford assay and samples were equalized by dilution with an appropriate volume of KPBS in Low-Bind Eppendorf tubes. Samples were then centrifuged 7500 × g for 15 min in 4 °C and pellets were thoroughly resuspended in the original volume of KPBS (3–5× original pellet volume) to gain total membrane fraction (TMF). We verified that 7500 × g is sufficient to pellet most of the LAMP2/TMEM192-positive fraction while preventing excessive aggregation of membranes (**Figure S3 a**). TMF were next incubated with 40 μL prewashed anti-HA magnetic bead slurry (Pierce) for 30 min in 4 °C. This was followed by two washes with KPBS supplemented with 150 mM NaCl, a transfer to a clean tube, and three additional washes with KPBS. Captured lysosomal proteins were then eluted with 100 µL of 0.5% NP-40 in KPBS without protease inhibitors (30 min, 4 °C), separated from beads and snap-frozen. Beads were then boiled in 100 µL of 1× reducing buffer and (“bead boil”) to check elution quality. Efficiency of lysosome purification was verified for each replicate by Western blotting a 10% SDS PAGE gel and staining against a panel of lysosomal (LAMP2, TMEM192, RAB7, BORCS6) and nonlysosomal (EEA1, BiP, Calnexin) proteins (**Figure S3 b**).

### Mass spectrometry

Sample volume was adjusted to 100 µL with 1% deoxycholate (DOC) solution in 100 mM ammonium bicarbonate (ABC) buffer, pH 8.0. Proteins were purified by adding 400 µL acetonitrile and centrifugation at 16,000 g for 5 min. The pellet was washed 2× with 300 µL 80% ethanol and 1× with 300 µL 80% methanol, dried in a vacuum concentrator, and finally dissolved in 50 µL 100 mM ABC buffer, pH 8.0. Proteins were reduced by adding 50 µL 10 mM DTT solution at 56°C for 30 min and alkylated with 50 µL 550 mM IAA solution at room temperature for 20 min and digested overnight at 37°C by adding 0.5 µg trypsin (Promega).

40% of the digested samples were analyzed using an UltiMate 3000 nano-HPLC system coupled to an Orbitrap Eclipse mass spectrometer (Thermo Scientific, Bremen, Germany). The peptides were separated on a homemade frit-less fused-silica micro-capillary column (75 µm i.d. x 280 µm o.d. x 16 cm length) packed with 2.4 µm reversed-phase C18 material (Reprosil). Solvents for HPLC were 0.1% formic acid (solvent A) and 0.1% formic acid in 85% acetonitrile (solvent B). The gradient profile was as follows: 0–4 min, 4% B; 4–117 min, 4–30% B; 117–122 min, 30–100% B, and 122–127 min, 100% B. The flow rate was 250 nL/min.

The Orbitrap Eclipse mass spectrometer equipped with a field asymmetric ion mobility spectrometer (FAIMS) interface operated in the data dependent mode with compensation voltages (CV) of -45, -55 and -75 and a cycle time of one second. Survey full scan MS spectra were acquired from 375 to 1500 m/z at a resolution of 60000 with an isolation window of 1.2 mass-to-charge ratio (m/z), a maximum injection time (IT) of 118 ms, and automatic gain control (AGC) target 400,000. The MS2 spectra were measured in the Orbitrap analyzer at a resolution of 15,000 with a maximum IT of 22 ms, and AGC target or 50,000. The selected isotope patterns were fragmented by higher-energy collisional dissociation with normalized collision energy of 30%.

### Analysis of MS data

Data analysis was performed using Proteome Discoverer 3.1 (Thermo Scientific) with search engine Sequest. The raw files were searched against Uniprot *Homo sapiens* database. Precursor and fragment mass tolerance was set to 10 ppm and 0.02 Da, respectively, and up to two missed cleavages were allowed. Carbamidomethylation of cysteine was set as static modification, oxidation of methionine was set as a variable modification. Peptide identifications were filtered at 1% false discovery rate. Protein abundance was normalized to the total peptide amount. Protein abundance ratio between BicD2 and KIF5C samples was calculated as a median of all peptide abundance ratios for each replicate and then a final median value was calculated for 3 replicates. Z-score-based P-value was calculated by comparing the observed protein log_2_ abundance ratio to a global background distribution of log_2_ ratios and corrected for multiple testing by the Benjamini-Hochberg method.

R was used to evaluate replicate similarity using the Pearson correlation. The statistic was then visualized as a heatmap by distance on a correlation plot.

To control for variance reliability in low peptide count samples, DEqMS (Zhu et al., 2020) was used to model mean-variance relationship. Three comparisons were used with the grouping structures of BicD2 (+A/C, ctrl), KIF5C (+AC, ctrl), stimulated samples (KIF5C + A/C, BicD2 + A/C). The matrix model used for the three comparisons was the same (∼0 + group) denoting each coefficient was independently calculated. Finally, for the control vs. stimulated contrast, we used the stimulated data as the numerator, and the control as the denominator. For the comparison of the two stimulated conditions, KIF5C was used as the numerator, and BicD2 was used as the denominator. The data for each comparison were plotted with the mean-variance model to verify the required relationship between variance and peptide count to meet the priors of the DEqMS analysis. Contrasts that failed to meet priors due to inconsistent relationship were evaluated using the Limma model adjusted P-value and limma z-score for downstream analysis. Adjusted P-values and fold-change values produced by the appropriate correction were used for volcano plots and enrichment analysis.

Proteins were deemed enriched against the uninduced control (background) if they complied with one of the following criteria: 1) protein in at least one of the stimulated samples was significantly enriched against the background control (DEqMS adj.P-val<0.05); or 2) protein was similarly enriched using the Protein Discoverer criteria as a threshold (adj.P.val<0.05) and was detected in all three replicates, with at least 2 peptides used for the quantification.

### Enrichment analysis

For subcellular compartments, proteins enriched against the background were assigned to selected organelles (lysosome, endosome, plasma membrane, mitochondrion, peroxisome, endoplasmic reticulum, Golgi apparatus, cytosol) using the COMPARTMENTS (human, integrated) dataset (Binder et al., 2014) downloaded from https://compartments.jensenlab.org (11/2025). Only assignments with confidence score≥4 were filtered. Enrichment of each organelle in the MS dataset was calculated as a ratio of all human proteins assigned to a given organelle and a number of such proteins identified in the MS dataset. To test whether ER/Golgi proteins showed a directional enrichment between samples BicD2 and KIF5C, we performed logistic regression using group membership (binary) as the response variable and log₂ fold change (KIF5C/BicD2) as the predictor. Statistical significance and directionality of the association were assessed using a binomial generalized linear model. Log₂ fold change distribution of ER/Golgi and non-ER/Golgi proteins was plotted on a split violin plot with R (ggplot2, geom_split_violin).

For FGSEA, we used the z-score metric generated from the appropriate variance correction model to build our ranked universe. We then converted our entrez ID to gene symbols and removed any replicated hit. Specific gene sets were downloaded from the molecular signature database (MsigDB) (Subramanian et al., 2005). Clusterprofiler (Yu et al., 2012) and Fast Gene set Enrichment (Korotkevich et al., 2021) packages were used. Gene sets with at least 10 and at most 500 genes were included in the enrichment analysis.

### SDS PAGE and Western blotting

Samples for SDS polyacrylamide gel electrophoresis were made by washing cells 2× in PBS and scraping them into a suitable volume of lysis buffer (50 mM Tris pH 7.5, 150 mM NaCl, 1% Triton X-100, 10% glycerol, 0.5 mM EGTA, 0.5 mM EDTA, 5 mM Na_2_P_2_O_7_, 50 mM NaF, 1 mM sodium orthovanadate, 10 μg/mL aprotinin, 1 μg/mL pepstatin, 10 μg/mL leupeptin, 0.4 mM Pefablock SC). After clearing lysates by centrifugation at 16,000 × g for 10 min, Bradford Plus Protein Assay Reagent (Thermo Fisher) was used to equalize protein concentration across samples. Samples were then denatured by adding 5× sample buffer (40 mM Tris-HCl pH 6.8, 5% glycerol, 1% SDS, 100 mM DTT, 0.0025% bromophenol blue) and incubating at 95 °C for 10 min. Samples were separated by SDS PAGE (Bio-Rad apparatus) and transferred to nitrocellulose membranes (Amersham Protran Premium 0.2 NC, Cytiva) using a wet blotting procedure (25 mM Tris, 192 mM glycine, 20% methanol). Membranes were blocked in 5% milk in TBS-T (20 mM Tris-HCl pH 7.6, 137 mM NaCl, 0.05% Tween-20) and incubated overnight with primary antibodies. After washing (3× 5 min with TBS-T), membranes were incubated with HRP-conjugated secondary antibodies, washed again, and chemiluminescence was detected with an ECL substrate (WesternBright, Biozym) on a Fusion FX imaging system (Vilber). Densitometry from digital images was accomplished with ImageJ (Analyze > Gels); raw numbers were first normalized to a sample loading control and then to an untreated (control) sample.

## Supplementary data

- Video 1
- Table 1
- Table 2
- Supplementary Figure 1
- Supplementary Figure 2
- Supplementary Figure 3
- Supplementary Figure 4

## Acknowledgements and funding sources

The authors thank the Biooptics and the Protein core facilities from the Medical University of Innsbruck for their support.

This research was funded in whole or in part by the Austrian Science Fund (FWF) (DOI 10.55776/P32608, DOI 10.55776/P36975 and DOI 10.55776/P35805B). For open access purposes, the author has applied a CC BY public copyright license to any author accepted manuscript version arising from this submission.

## Data availability

Relevant code is available at github.com/VojtechDostal/imagej-scripts (microscopic analysis) and https://github.com/arpollio/lysosomal_repositioning_proteomics (proteomic analysis). The mass spectrometry proteomics data have been deposited to the ProteomeXchange Consortium via the PRIDE partner repository with the dataset identifier PXD075882. Reviewer access details: **Project accession:** PXD075882. **Token:** GriyRczVY0u7. Vectors will be deposited at Addgene (https://addgene.org/Lukas_Huber).

---

**Video 1**

Time-lapse confocal microscopy of U2OS repositioning cell lines showing the temporal dynamics of lysosomal relocalization into the perinuclear (left) or peripheral area (right) upon 5nM A/C treatment. Overlay of mCherry (green) with phase contrast (grey), 1 frame/5 min; see also **Figure 1 e**.

**Figure S1.**
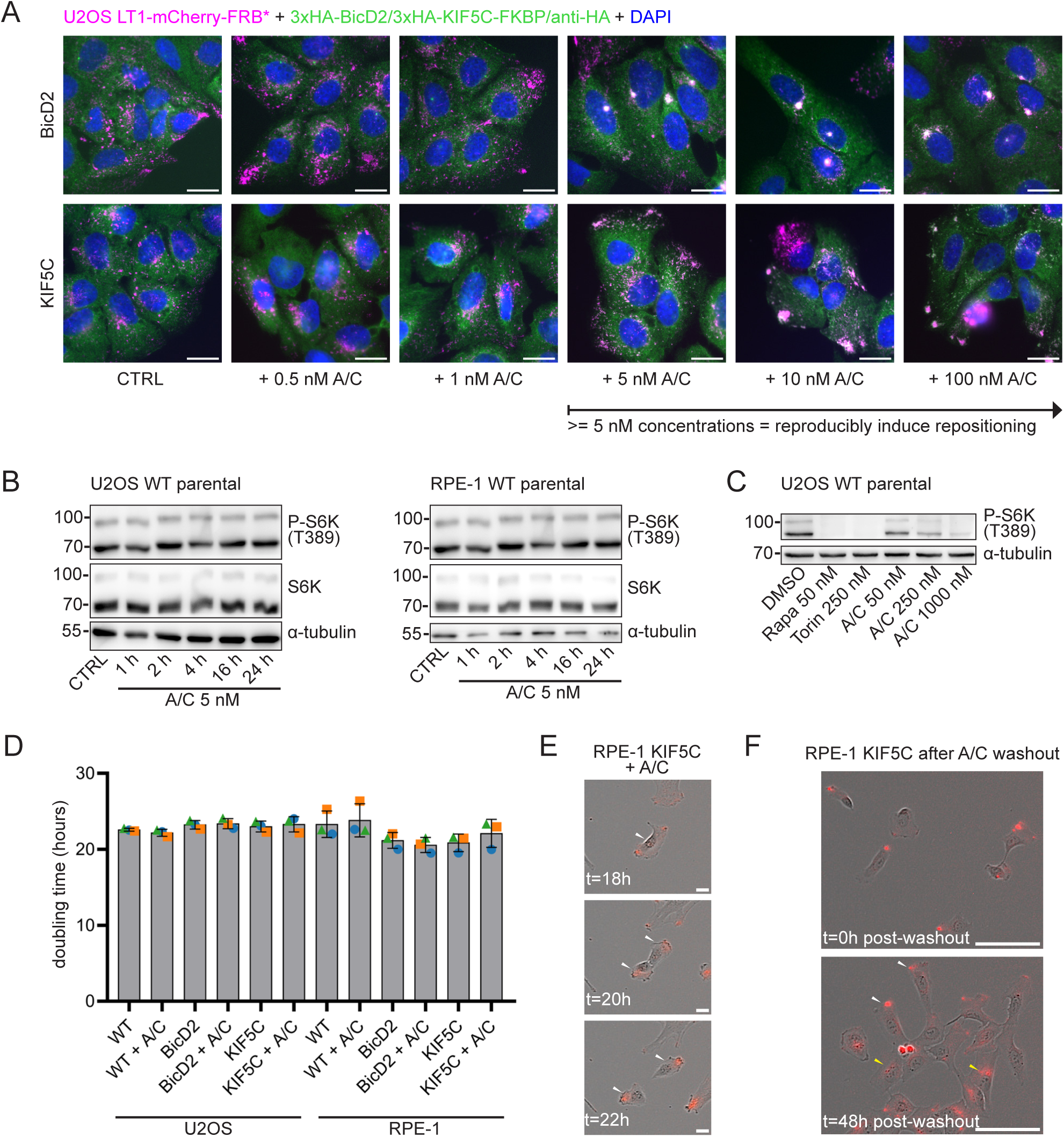
**A** Immunofluorescence staining of U2OS repositioning cell lines induced for 30 min with indicated concentrations of A/C. Magenta: mCherry (LT1-mCherry-FRB*), green: HA (3×HA-BicD2/KIF5C-FKBP), blue: DAPI (nuclei). Scale bar: 20 μm. **B** Immuno-blotting of cell lysates from U2OS (left) and RPE-1 (right) parental cells repositioned with 5 nM A/C for 1–24 h, showing lack of effect on mTORC1 signalling. **C** Immuno-blotting of cell lysates from U2OS parental cells treated with 50 nM rapamycin, 250 nM torin or high concentrations of A/C where mTORC1 inhibition is observed. **D** Doubling time of parental and repositioning U2OS and RPE-1 cell lines with or without A/C. N=3 independent experiments. Error bars: mean ± SD. **E** DIC and mCherry (red) overlay image of RPE-1 KIF5C cell line (Incucyte). Cells divide and maintain the lysosomal localization even during division (arrows point to daughter cells). Scale bar: 20 μm. **F** DIC and mCherry (red) overlay image of RPE-1 KIF5C cell line. Repositioning effect gradually dissipates after removal of A/C from the medium. At 48 hours approximately 50% of cells were repositioned (white arrows), the rest had random lysosomal localization (yellow arrows). Scale bar: 100 μm.

**Figure S2.**
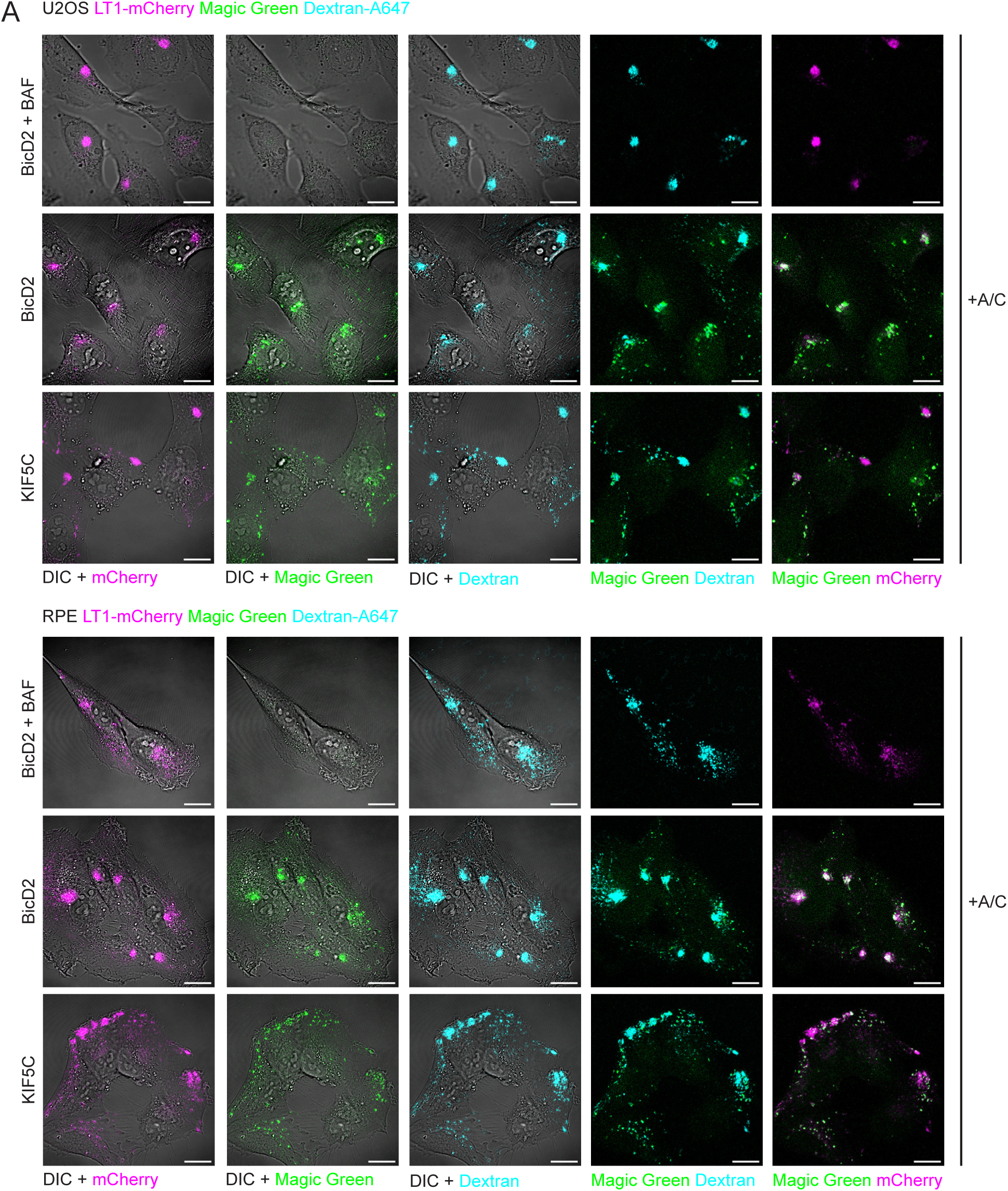
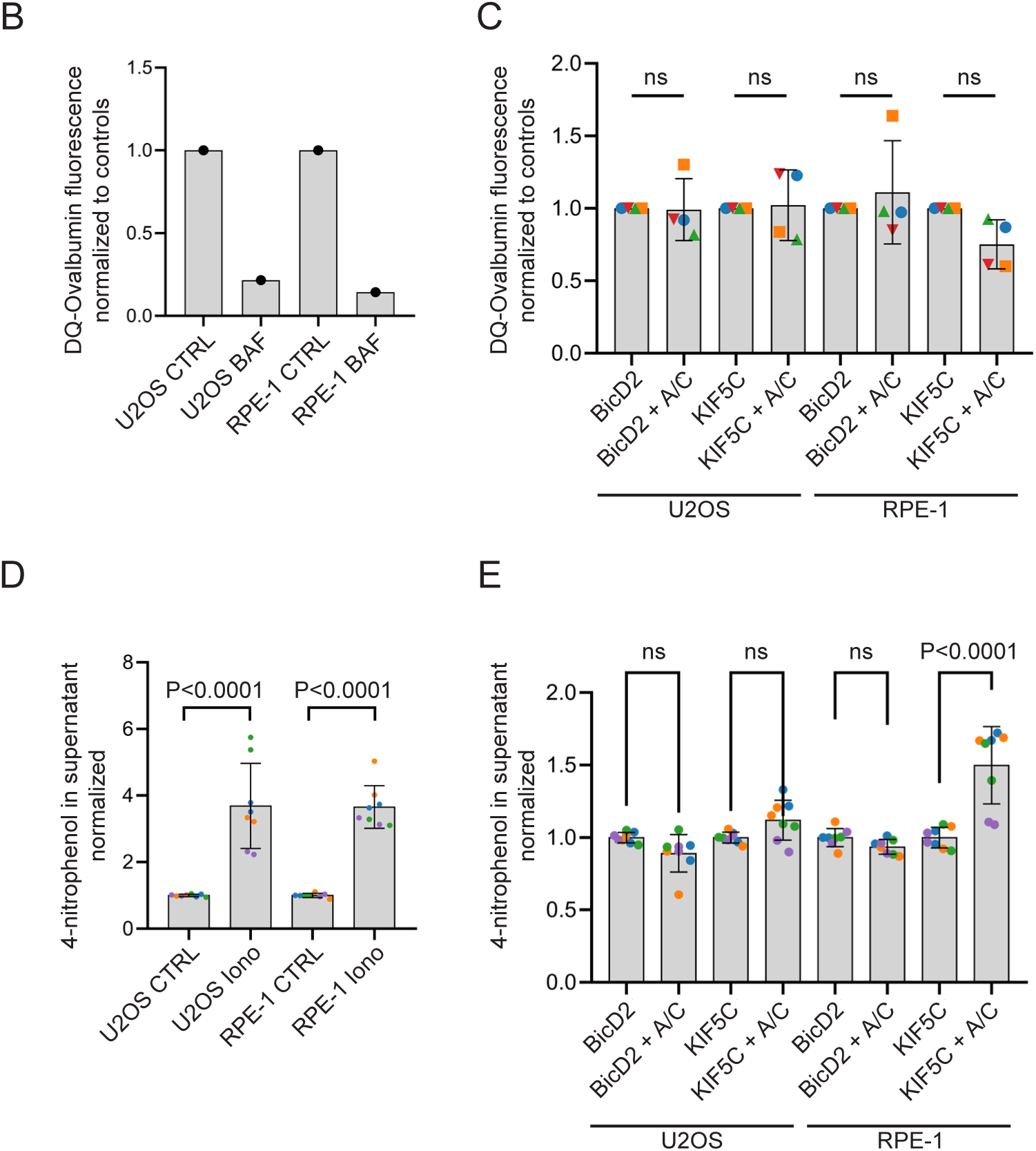
**A** Degradative capacity of lysosomes of U2OS and RPE-1 cell lines measured with a Magic Green assay, normalized via Dextran-Alexa 647 and controlled with a bafilomycin (BAF)-treated sample. 200 nM BAF was added 2h before Magic Green treatment; decrease in fluorescence in BAF-treated cells confirms specificity of the Magic Green signal for lysosomal enzymatic activity. Scale bar: 15 μm. **B** Plate reader assay of total DQ-Ovalbumin fluorescence in U2OS and RPE-1 cells. 200 nM BAF (added 2 h before DQ-Ovalbumin treatment) reduces the production of the fluorescent product, indicating assay specificity for lysosomal proteolytic activity. N=1 biological replicate. **C** Quantitative plate reader assay of DQ-Ovalbumin fluorescence in U2OS and RPE-1 repositioning cell lines. Total proteolytic activity is not significantly altered in repositioned cell lines. N=4 biological replicates. Test: Ordinary one-way ANOVA. Error bars: Mean ± SD. **D** Plate reader assay for β-hexosaminidase (BHEX) activity in the medium of control and 2 μM io-nomycin-treated U2OS and RPE-1 cells. N=4 biological replicates. Test: Ordinary one-way ANOVA. Error bars: Mean ± SD. **E** Plate reader assay for BHEX activity in the medium of U2OS and RPE-1 repositioning cell lines, normalized to total intracellular BHEX activity. N=4 biological replicates. Test: Ordinary one-way ANOVA. Error bars: Mean ± SD.

**Figure S3.**
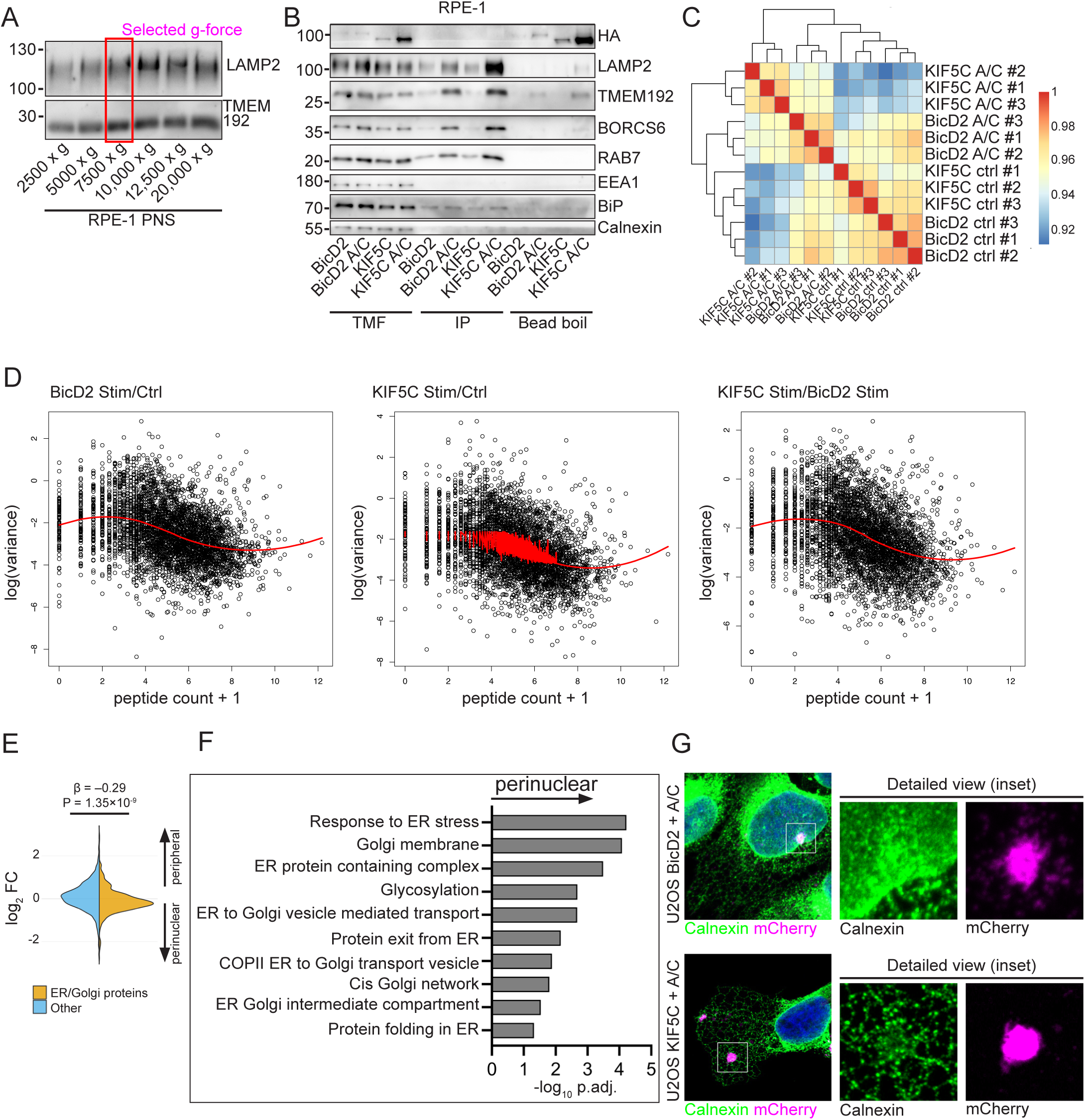
**A** Western blotting analysis of post-nuclear supernatant (PNS) from RPE-1 WT cells centrifuged at different speeds revealed that a majority of the lysosomal fraction pellets at 7,500 x g, selected for subsequent experiments. **B** Western blotting analysis of input fraction, elutions and non-eluted proteins on beads from a representative LysoIP experiment with BicD2 and KIF5C RPE-1 cell lines. Note that the anti-HA membrane was assembled from two previously cut pieces of membrane, producing a weak line at the cut site. TMF = total membrane fraction, IP = eluted fraction. **C** Crosswise QC comparison of Pearson correlation values. Note that the control samples cluster together showing consistent correlation before treatment. **D** Fitted mean-variance plots denoting the relationship between variance and peptide count. The two outer plots show a normal relationship where peptide count and variance are related, thus meeting the priors required for DEqMS. The central plot fails to meet the priors due to the inconsistent shape in the middle peptide counts denoting an inconsistent relationship. For this central comparison the p_adj_ for the less conservative limma model output was used instead of the DEqMS. **E-G** LysoIP for the quantification of ER-lysosome / ER-Golgi contacts. **E** Split violin plot showing proteins detected in LysoIP and annotated as components of ER or Golgi (yellow) or other proteins (blue). Statistical test: Logistic regression. **F** Enriched perinuclear GO gene sets sorted by p-value specifically denoting ER and Golgi function. **G** Airyscan super-resolution staining of endoplasmic reticulum (calnexin, green) in U2OS cells, showing organization of the ER tubules around peripheral lysosomal patches.

**Figure S4.**
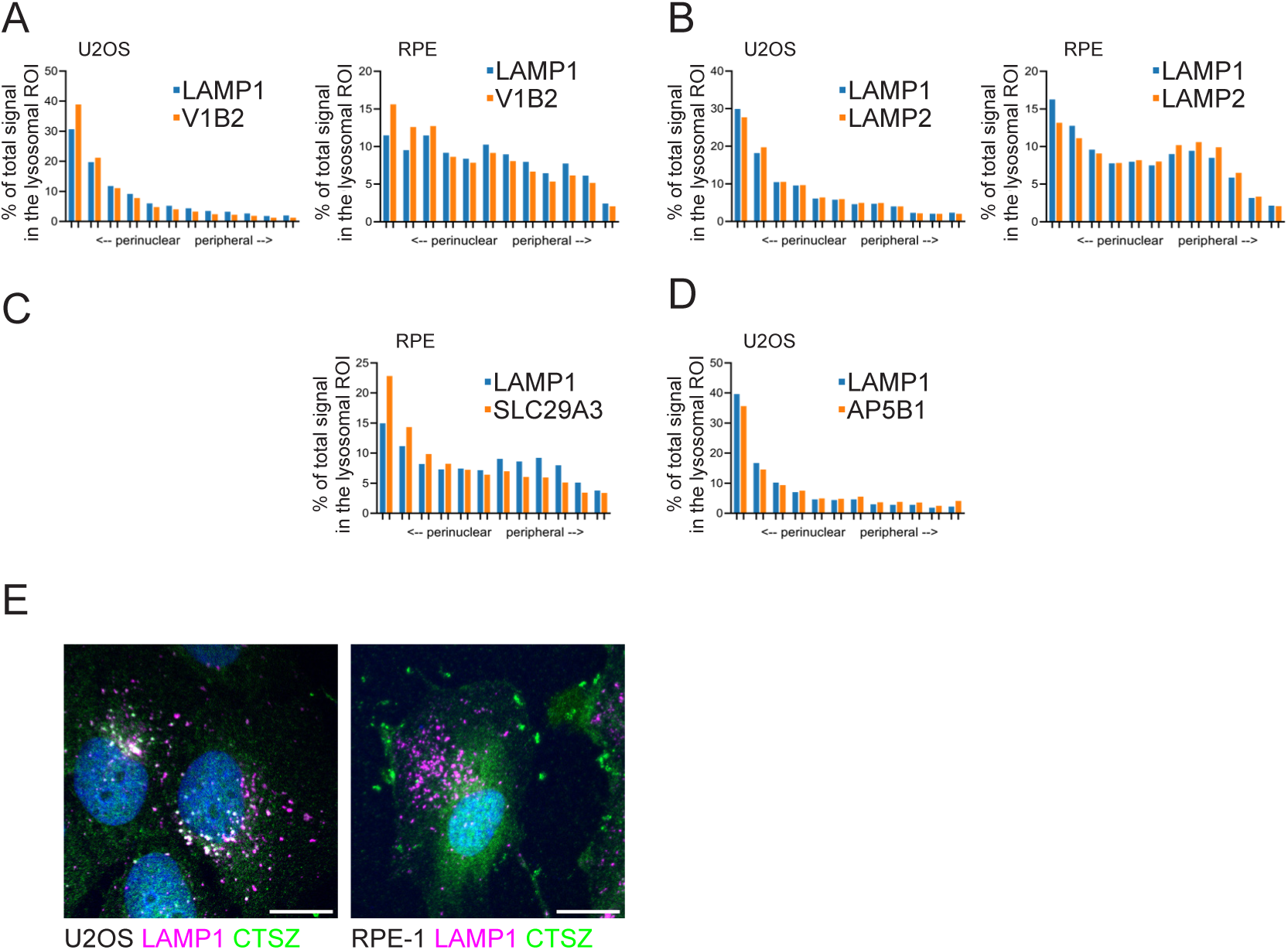
**A-D** Histograms of signal intensity in 12 equally sized areas of increasing distance from the nucleus (1=most perinuclear, 12=most peripheral). % of total signal in each area was plotted here (validated hit was not normalized to LAMP1), showing distinct intensity profiles of lysosomes in U2OS and RPE cells. **E** Cathepsin Z (CTSZ) is a candidate for peripheral lysosomes in RPE, and shows diffuse distribution and localization to patches on the plasma membrane on immunofluorescence; CTSZ in U2OS has a perinuclear distribution on immunofluorescence.

